# A *trans-*splicing model for poly(A)-tail addition during VSG mRNA maturation in *Trypanosoma brucei*

**DOI:** 10.64898/2026.10.18.760011

**Authors:** Nikolay G. Kolev, Huafang Shi, Christian Tschudi

**Affiliations:** Department of Epidemiology of Microbial Diseases, Yale School of Public Health, New Haven, CT, USA

## Abstract

Trypanosomes are characterized by polycistronic transcription and a pre-mRNA processing pathway in which spliced leader (SL) *trans*-splicing and cleavage and polyadenylation produce individual mature mRNAs. SL and poly(A)-tail addition within the intergenic regions are obligatorily coupled, yet the molecular basis for this coupling has remained unknown. Here we identified a splicing enhancer sequence required for efficient maturation of the highly abundant variant surface glycoprotein (VSG) mRNA and showed that it recruits factors implicated in RNA processing and nuclear export, including the conserved RNA helicase Sub2. RIP-seq revealed that Sub2 associates not only with VSG pre-mRNA but also with SL RNA precursors carrying long genome-encoded 3′ extensions. Integration of these findings with analyses of published U1C iCLIP and cold shock domain protein 1 (CSD1) seCLIP datasets supports a model in which polyadenylated SL RNA precursors, rather than mature SL RNPs, serve as substrates for *trans*-splicing. We propose that VSG pre-mRNA forms an integral component of a specialized spliceosomal assembly that catalyzes successive rounds of SL and poly(A)-tail *trans*-splicing, driven by extensive base-pairing interactions. This model provides a mechanistic framework for the strong coupling of mRNA 5′- and 3′-end formation in trypanosomes and suggests that spliceosome-catalyzed polyadenylation represents an evolutionarily ancient strategy for eukaryotic pre-mRNA processing. More broadly, it challenges the notion that the two transesterification steps of RNA splicing require a universally conserved arrangement of spliceosomal RNA components.

## INTRODUCTION

*Trypanosoma brucei*, an early-diverging single-celled eukaryotic parasite and the causative agent of African trypanosomiasis, expresses its protein-coding genes as long polycistronic transcripts that are processed into individual mRNAs. This maturation relies on spliced leader (SL) *trans*-splicing,^1,2^ which adds a capped 39-nt SL sequence to the 5′ end of every mRNA, coupled spatially ^3–5^ and functionally ^6,7^ to cleavage and polyadenylation upstream of the *trans*-splice site. Despite conservation of the cleavage and polyadenylation specificity factor (CPSF) complex,^8^ including the endonuclease CPSF73, pre-mRNAs in *T. brucei* lack a defined polyadenylation signal analogous to the canonical AAUAAA motif in other eukaryotes. Consequently, the mechanisms that specify cleavage sites and coordinate SL *trans*-splicing with polyadenylation remain poorly understood.

A mechanistic model for SL *trans*-splicing in trypanosomes proposed in the 1980s ^9,10^ posits a two-step transesterification reaction analogous to *cis*-splicing (Figure 1A). However, experimental support for this model remains limited beyond the identification of a Y-structure splicing intermediate.^9,10^ Notably, several key features of canonical spliceosome function are absent or divergent in trypanosomes. For example, U2 snRNA lacks the conserved branch-site recognition sequence,^11–15^ and branch-point adenosines in pre-mRNAs have been difficult to define.^16,17^ The role of U1 snRNA in *trans*-splicing also remains unclear.^18^ In addition, SL RNA is assembled into Sm-containing ribonucleoprotein particle prior to *trans*-splicing.^19,20^ Sm snRNPs are known to be extremely stable,^21^ raising questions about SL RNP Sm components turnover and reuse. Together, these observations suggest that SL *trans*-splicing in trypanosomes may proceed through a mechanism that differs substantially from canonical spliceosome-mediated pre-mRNA *cis*-splicing. SL RNA itself is extensively modified, carrying a hypermethylated cap structure (cap 4)^22^ and a pseudouridine at position 28 (Ψ28).^23^ While the co-transcriptionally obtained cap 4 modifications are transferred to all mature mRNAs,^24,25^ pseudouridylation has not been reported for mRNA-attached SL sequences, challenging predictions that SL Ψ28 will be ubiquitous in *T. brucei* mRNA.^23^ These discrepancies suggest that the substrates and intermediates of SL *trans*-splicing in trypanosomes remain incompletely characterized. Moreover, the unusual features of trypanosome spliceosomal RNAs raise the possibility that the two transesterification reactions of RNA splicing can proceed through molecular interactions distinct from those involved in canonical *cis*-splicing.

**Figure 1.**
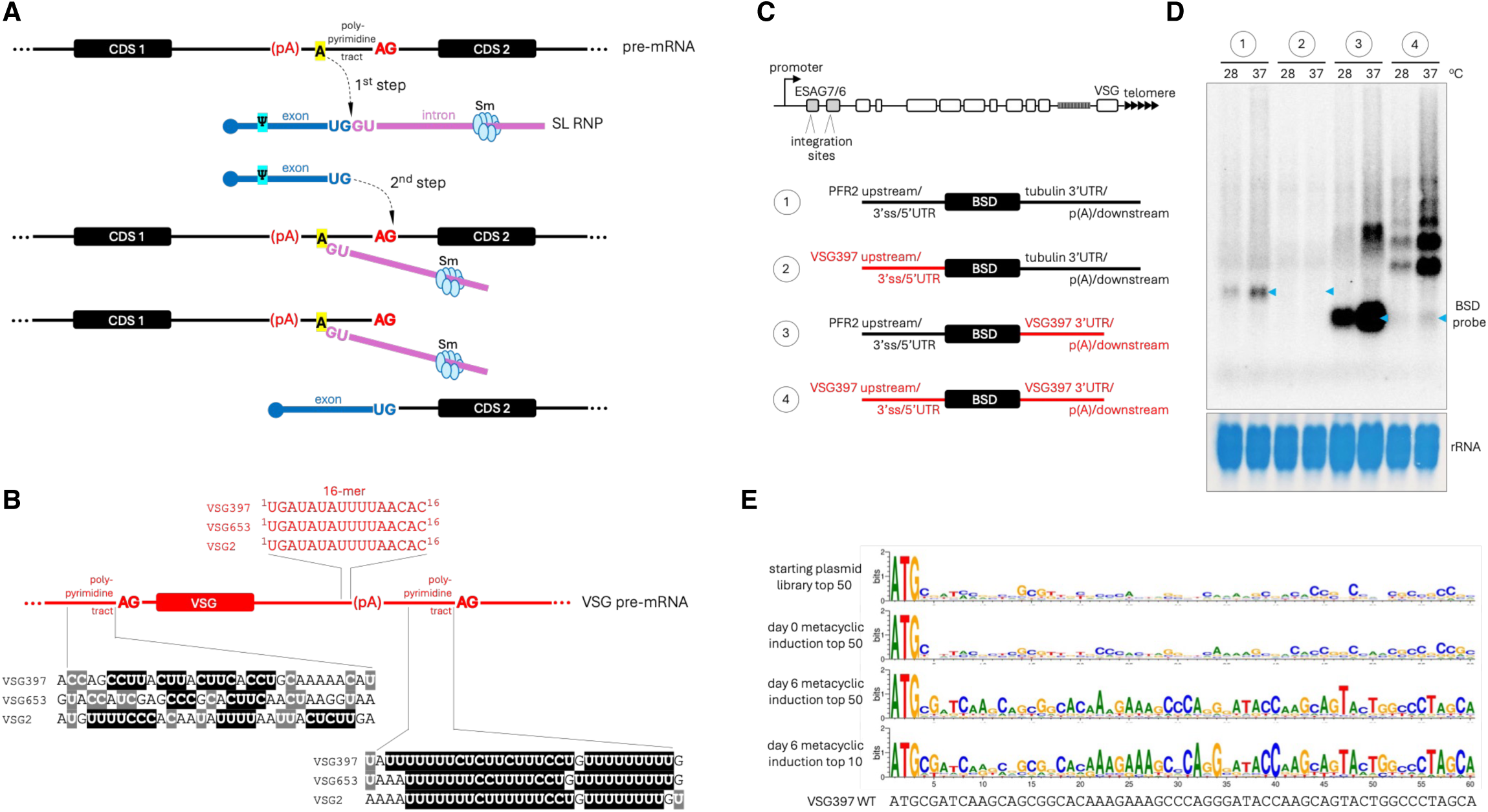
VSG pre-mRNA has a weak SL *trans*-splice site. (A) Classical model of SL *trans*-splicing in trypanosomes proposed in the 1980s. (B) Sequences of the conserved 16-mer and the upstream and downstream polypyrimidine tracts in VSG pre-mRNA. (C) Reporter constructs used to assess the strength of the VSG 3′ splice site for SL addition. A schematic representation of a bloodstream-form VSG expression site indicates the possible integration sites. (D) Northern blot analysis of BSD reporter mRNA abundance in procyclic cells cultured for 4 h at the indicated temperatures. Blue arrowheads indicate the expected size of properly matured mRNAs for each construct. (E) Mutagenized library screen used to identify sequence elements within the first 60 nt of the VSG397 CDS required for efficient mRNA production.

The variant surface glycoprotein (VSG) mRNA, which accounts for ∼10% of total mRNA in bloodstream forms (BF, in the mammalian host) and metacyclic forms (MF, in the insect vector), provides a powerful model to investigate these mechanisms. VSG is essential for infectivity and survival in mammalian tissues harboring the parasite and is one of the best studied examples of antigenic variation.^26,27^ VSG genes exhibit monogenic expression from specialized telomeric Pol I transcription units termed metacyclic and bloodstream expression sites (MES and BES, which are monocistronic and polycistronic, respectively). The active ES is found in a Pol I expression site body^28,29^ and is positioned in proximity to one of the SL RNA multi-copy gene array loci,^30,31^ highlighting the importance of access to large amounts of SL RNA substrate for the high-level expression of VSG mRNA. Despite its high abundance, the VSG pre-mRNA has a suboptimal polypyrimidine tract containing fewer uridines and multiple interrupting purines at the SL addition site, in contrast to other highly expressed *T. brucei* transcripts. Conversely, a strong polypyrimidine tract is located downstream of the polyadenylation site but is not used for *trans*-splicing (Figure 1B), as SL *trans*-spliced sequences corresponding to this position are nearly absent in available RNA-seq data. In addition, a highly conserved 16-nt sequence element upstream of the polyadenylation site is required for high-level VSG expression, although its function remains unknown.^32–34^ These features suggest that VSG pre-mRNA processing relies on non-canonical sequence determinants.

Here, we identified a splicing enhancer at the 5′ end of the VSG coding sequence that is required for efficient pre-mRNA processing and recruits factors involved in RNA processing and nuclear export. We showed that this element has the potential to be involved in an extensive RNA-RNA interaction with a downstream sequence in the VSG pre-mRNA and that it associates with factors known to facilitate dynamic intermolecular base-pairing contacts during RNA editing in the mitochondrion. Based on these findings, we propose a model in which the conserved 16-mer acts as a central organizing element in a spliceosomal base-pairing network that coordinates SL *trans*-splicing and polyadenylation. In this model, both the SL sequence and the poly(A) tail are transferred from a polyadenylated SL RNA precursor through successive *trans*-splicing reactions, providing a potentially general mechanism for mRNA maturation in trypanosomes. Furthermore, our model suggests that the conserved chemistry of RNA splicing can be supported by distinct molecular architectures, revealing previously unappreciated mechanistic diversity in eukaryotic pre-mRNA processing.

## RESULTS

### VSG397 pre-mRNA has a weak SL *trans*-splice site that requires a splicing enhancer

The highly degenerate polypyrimidine tract at the SL addition site in VSG pre-mRNA (Figure 1B) deviates substantially from optimal highly functioning sequences for pre-mRNA splicing, which are typically 15 to 20 nt long and contain continuous stretches of uridines. Purine nucleotides (A or G) are generally excluded, as their presence significantly weakens spliceosome binding. This suggests the existence of additional sequence elements required for the efficient maturation of the most abundant mRNA in *T. brucei*. To determine if the VSG SL *trans*-splice site is sufficient on its own for proper mRNA maturation, we used a series of reporter constructs carrying the blasticidin-S deaminase (BSD) coding sequence (Figures 1C and S1A). The upstream sequences were either from the paraflagellar rod protein 2 (PFR2) intergenic region containing the PFR2 SL *trans*-splice site or 508 nt upstream of the VSG397 coding sequence (CDS). The downstream sequences, containing polyadenylation sites, were the β/α-tubulin intergenic region or the VSG397 sequence extending from the CDS stop codon to the downstream non-functional SL splice site, which is preceded by an optimal (Figure 1B) polypyrimidine tract. The reporter constructs were targeted for integration into the CDS of ESAG7/6 (Expression-Site-Associated Genes 7 and 6, encoding the transferrin receptor heterodimer) at the promoter-proximal region of the Pol I-transcribed bloodstream VSG expression sites (Figure 1C) and were individually transfected into procyclic (insect-form) cells. We chose procyclic cells to minimize any effects of 5′UTR and 3′UTR sequences on mRNA stability, as the VSG mRNA-specific stabilization factor cyclin F-box protein 2 (CFB2)^34,35^ is expressed exclusively in bloodstream form cells, and PFR2 and tubulin mRNAs are constitutively abundant in both procyclic and bloodstream-form cells. The promoter-proximal region of bloodstream VSG expression sites was selected as the integration site because Pol I initiation and transcription occur in these regions even in procyclic cells,^36,37^ and transcriptional activity can be further increased by elevating the temperature from 28°C to 37°C.^38^ Trypanosome populations carrying the integrated constructs were cultured at 28°C and 37°C, and total RNA was analyzed by northern blotting using a radiolabeled BSD probe (Figure 1D).

As expected, all detectable BSD mRNA species were 2- to 6-fold more abundant in the 37°C samples (Figure 1D). The reporter containing the VSG397 upstream sequence (construct 2) and the tubulin downstream sequence failed to produce detectable BSD mRNA (Figure 1D), indicating that the VSG upstream region containing the SL *trans*-splice site is not sufficient for detectable mRNA production. In contrast, the VSG397 downstream region in the absence of other VSG sequences (construct 3) supported robust production of properly matured mRNA of the expected size (Figure 1D). When the VSG397 downstream and upstream regions were combined in the same reporter (construct 4), BSD mRNA was detected at similar levels to those of construct 3, however, the larger size of the transcripts was consistent with 5′-end processing that occurred at the SL addition sites of ESAG7 and ESAG6, the genes used for integration (Figure 1D). Together with the absence of detectable mRNA from the construct containing only the upstream VSG sequence, these results confirmed that the VSG397 SL *trans*-splice site is intrinsically weak and indicated that additional sequence element(s) outside the tested VSG upstream and downstream regions are required for efficient SL addition at this site.

To determine whether the required sequences are located within the VSG397 CDS, we performed deletion analysis (Figure S2). Either 618 or 100 nt at the beginning of VSG397 CDS were sufficient to support expression of properly matured mRNA (Figure S2A), and the first 63 nt were more important than nt 64-99 (Figure S2B). To define the sequence requirements of the identified splicing enhancer at the beginning of the VSG397 CDS, we performed an *in vivo* screen using a doped mutagenesis library encompassing the first 60 nt of the VSG397 CDS (Figure S1C). Each position, with the exception of the start codon, was represented by doped nucleotides at a wild-type-to-mutant ratio of 7:3. Following selection of stable transformants, cells were induced to differentiate into metacyclic forms expressing the mutagenized VSG397 through inducible expression of RNA-binding protein 6 (RBP6).^39^ Expressed VSG397 transcripts were analyzed on day 6 of RBP6 induction by RT-PCR followed by next-generation sequencing, in parallel with PCR samples of the starting plasmid library and genomic DNA from the trypanosome population prior to induction (Figure 1E). Analysis of the expressed sequences failed to identify a discrete short element within CDS positions 4-60 at either time point. Instead, wild-type nucleotides were enriched throughout the splicing enhancer, with slightly stronger preference in the 3′ half of the element (Figure 1E). Thus, sequence identity across the entire VSG397 SL *trans*-splicing enhancer is important for its stimulatory effect on VSG397 mRNA production.

### Identification of proteins interacting with the VSG397 splicing enhancer

To isolate proteins interacting directly or indirectly with the splicing enhancer element, we used extracts from cells induced to express cold shock domain protein 1 (CSD1),^40^ as this protein, although not associated with the mature VSG mRNA,^40^ causes the most rapid known induction of metacyclic VSG gene expression in insect-form cells (Figure 2A).^40^ To minimize background noise in pulldown assays using the splicing enhancer RNA as bait, we tested several mutagenized shorter segments within the enhancer based on data from the *in vivo* library screen. Specifically, we selected sequences that were abundant on day 0 but absent on day 6 of induced metacyclic development (not shown). We ultimately chose a 15-nt wild-type RNA from the 3′ half of the splicing enhancer as bait and a mutant version of the same RNA, containing substitution that prevent VSG expression, as a control (Figures 2B and S2C). Proteins isolated from whole-cell extract by the wild-type and mutant biotinylated RNAs were identified by LC-MS/MS (Table S1 and Figure 2C). Excluding highly abundant translation factors, the top 10 proteins enriched in the wild-type RNA pulldown samples with functions in RNA metabolism included factors implicated in RNA transcription, processing, and nuclear export. We focused subsequent analyses on RBP16, TDP1, and Sub2 because RBP16 and TDP1 were the most highly enriched proteins, whereas Sub2 has established roles in mRNA biogenesis.

**Figure 2.**
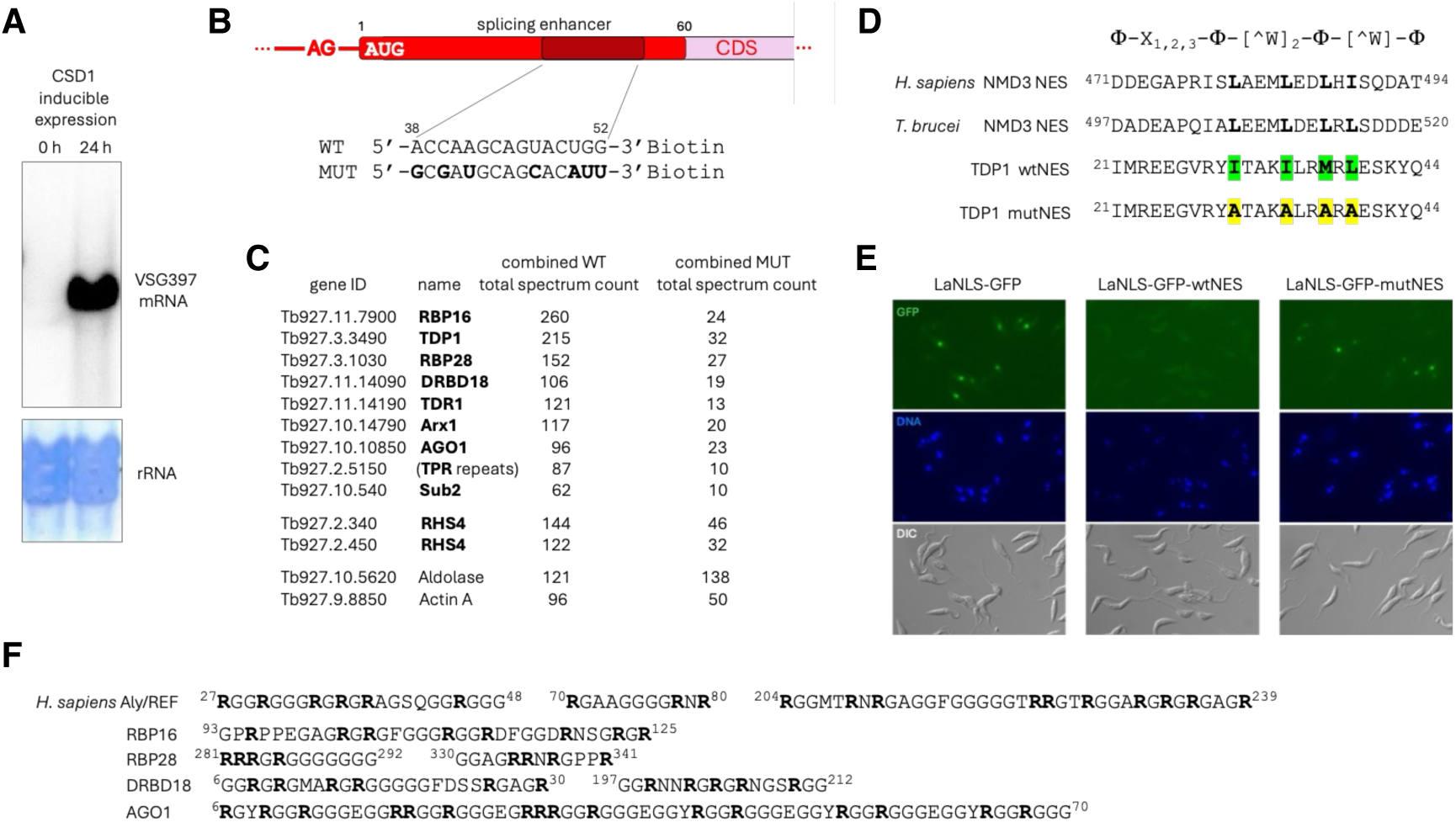
Identification of proteins interacting with the VSG397 splicing enhancer. (A) Northern blot analysis of VSG397 mRNA abundance in cells expressing CSD1. (B) Sequences of the 15-nt wild-type and mutant RNA baits used in the pulldown assay. (C) Top 10 proteins involved in RNA metabolism identified by LC-MS/MS as enriched in pulldowns with the wild-type RNA bait. Shown is the total spectrum count across all experiments. Aldolase and Actin A are included for comparison. (D) Nuclear export signal (NES) sequences from human and trypanosome NMD3 and the putative NES of TDP1. The consensus sequence for this class of NES is shown above. Residues mutated to alanine are highlighted. (E) Live-cell imaging of motile cells stably transfected with the indicated GFP constructs. (F) RG-rich regions in human Aly/REF and several of the proteins identified in the pulldown assay.

The proteins identified in the pulldown assay strongly emphasized the intricate functional connections between pre-mRNA processing and nuclear export. In addition to the TREX (TRanscription-EXport) complex component Sub2, the pulldown recovered several proteins with established or predicted roles in mRNP export. The high-mobility group protein TDP1 is a potent positive regulator of VSG expression that associates with active VSG expression site chromatin and is enriched in both the nucleolus and the Pol I VSG expression site body.^41^ Inspection of the TDP1 sequence revealed a potential class 1a nuclear export signal (NES)^42^ for binding to exportin 1/chromosome region maintenance 1 (XPO1/CRM1) at its N-terminus (Figure 2D). A similar class 1a NES consisting of four hydrophobic residues, with characteristic spacing lacking tryptophan, is present in the *T. brucei* NMD3 (Figure 2E), which functions in nuclear export of rRNA and Pol I-transcribed mRNAs.^43^ To determine whether the TDP1 NES is functional, we used a localization assay based on GFP fusions containing the nuclear localization signal (NLS) of the *T. brucei* La protein.^44^ Whereas LaNLS-GFP localized to the nucleus, fusion of TDP1 amino acids 21-44 to the GFP C terminus (LaNLS-GFP-wtNES) prevented nuclear accumulation (Figure 2E). A mutant version of the TDP1 peptide in which the hydrophobic NES residues were replaced with alanines (LaNLS-GFP-mutNES) largely restored nuclear localization (Figure 2E), demonstrating that the predicted NES in TDP1 is functional. Although inhibition of XPO1/CRM1 with leptomycin B was inconclusive because of trypanosome toxicity, our data support a role for TDP1 as a nuclear export adaptor that mediates XPO1/CRM1-dependent RNA export to the cytoplasm.

Several additional proteins recovered in the pulldown assay further linked RNA processing to nuclear export. Arx1 is an inactive methionine aminopeptidase that functions as a nuclear export receptor for the 60S ribosomal subunit in yeast.^45,46^ Retrotransposon hotspot proteins 4 (RHS4) in *T. brucei* are involved in pre-mRNA transcription and mRNA nuclear export, and their depletion results in the accumulation of polyadenylated RNA species lacking the SL in the nucleus.^47^ DRBD18 is an auxiliary factor in mitochondrial U-insertion/deletion RNA editing,^48^ but also functions in the nucleus, where it influences the selection of pre-mRNA SL-addition and polyadenylation sites^49,50^ and promotes nuclear export of a subset of mRNAs through interactions with Mex67/Mtr2.^51^ Trypanosomes lack an identifiable Aly/REF export factor,^52^ however, several of the proteins recovered in the pulldown contain RG-rich regions resembling those found in Aly/REF (Figure 2F). Among these proteins, the top hit, RBP16, is an auxiliary factor in mitochondrial U-insertion/deletion RNA editing^53^ that is also present in highly purified *T. brucei* nuclei.^54^ AGO1 is the trypanosome RNAi effector and functions in both the cytoplasm and nucleus,^55^ whereas RBP28 has no assigned function. As in Aly/REF,^56^ arginine methylation within the RG-rich regions is functionally important for both RBP16 and DRBD18.^57,58^ The function of Tudor domain protein 1 (TDR1) remains unknown, however, Tudor domains are known to recognize methylated arginine residues,^59^ suggesting a possible interaction between TDR1 and the methylated RG-rich regions of the identified proteins. Together, these findings suggest that the VSG397 splicing enhancer associates with a protein network that links pre-mRNA processing to assembly of export-competent mRNPs.

### Sub2 associates with VSG397 pre-mRNA and precursor SL RNA

Among the top proteins pulled down by the 15-nt segment of the splicing enhancer was the DEAD-box RNA helicase Sub2 (Figure 2C and Table S1), known in mammals as UAP56 (56-kDa U2AF^65^-associated protein) or DDX39B. It is the sole conserved subunit of the TREX complex in trypanosomes.^52^ Sub2 is an essential splicing factor^60^ with numerous proposed functions in pre-mRNA processing and mRNA nuclear export. It is required during the earliest stages of spliceosome assembly;^61^ it is critical for the U2 snRNP-branch-site interaction;^60^ contacts U4 and U6 snRNPs and can unwind the U4/U6 snRNA duplex;^61^ interacts directly with Aly/REF;^62,63^ and, acting as a molecular switch, orchestrates the transition of mRNA through several distinct molecular complexes from maturation to mRNP remodeling at the nuclear pore complex prior to export.^64^ In *T. brucei*, depletion of Sub2 by RNAi results in the accumulation of mRNA in the nucleus.^65^

To test whether Sub2 interacts with pre-mRNA *in vivo*, we C-terminally epitope-tagged one of its alleles and performed RIP-seq on immunoprecipitated transcripts from cells induced to differentiate into metacyclic forms. We also performed analogous experiments with the putative nuclear export adaptor TDP1 and with RBP16, the protein most strongly enriched in the splicing enhancer pulldown assays. RIP-seq analysis revealed that VSG pre-mRNA was abundant in the Sub2 immunoprecipitate despite being barely detectable in input RNA samples (Figures 3A and S3A). Reads containing the sequence immediately downstream of the 3′ splice-site AG dinucleotide indicated that 61.2 ± 2.4% (n=2) of the VSG397 RNA recovered in the Sub2 immunoprecipitate consisted of unspliced pre-mRNA lacking the SL. Compared with Sub2, TDP1 exhibited substantially lower enrichment for VSG pre-mRNA, with the exception of VSG639, an infrequently activated metacyclic VSG gene, for which precursor enrichment was comparable between TDP1 and Sub2 (Figures 3B and S3B). The lower level of VSG pre-mRNA association with TDP1 is consistent with its proposed role as an RNA nuclear export adaptor, as such factors would be expected to associate primarily with mature mRNPs competent for translocation through the nuclear pore complex. In addition, reciprocal co-immunoprecipitation experiments failed to detect an interaction between Sub2 and TDP1 (Figure S4), consistent with Sub2 dissociating from mature mRNPs prior to cargo loading and nuclear export.^64^ Under the experimental conditions used, RBP16 did not co-immunoprecipitate appreciable amounts of either VSG pre-mRNA or mature mRNA (Figures 3C and S3C). Nevertheless, RBP16 immunoprecipitates were enriched for the spliceosomal snRNAs U2, U4, U5 and U6 to higher levels (with the exception of U2) than Sub2 immunoprecipitates (Figure 3D). In contrast, TDP1 immunoprecipitates did not exhibit enrichment for spliceosomal snRNAs (Figure 3D). Together, these observations suggest that, unlike the prolonged association of Sub2 with pre-mRNA and mature mRNA during processing and mRNP remodeling before nuclear export, interactions between RBP16 and VSG (pre-)mRNA are likely to be transient and to occur during the catalytic stages of pre-mRNA splicing. This interpretation is supported by the similar enrichment of U2, U4, U5 and U6 in the RBP16 immunoprecipitates. The absence of spliceosomal snRNA enrichment in TDP1 immunoprecipitates is consistent with a role in RNA nuclear export.

**Figure 3.**
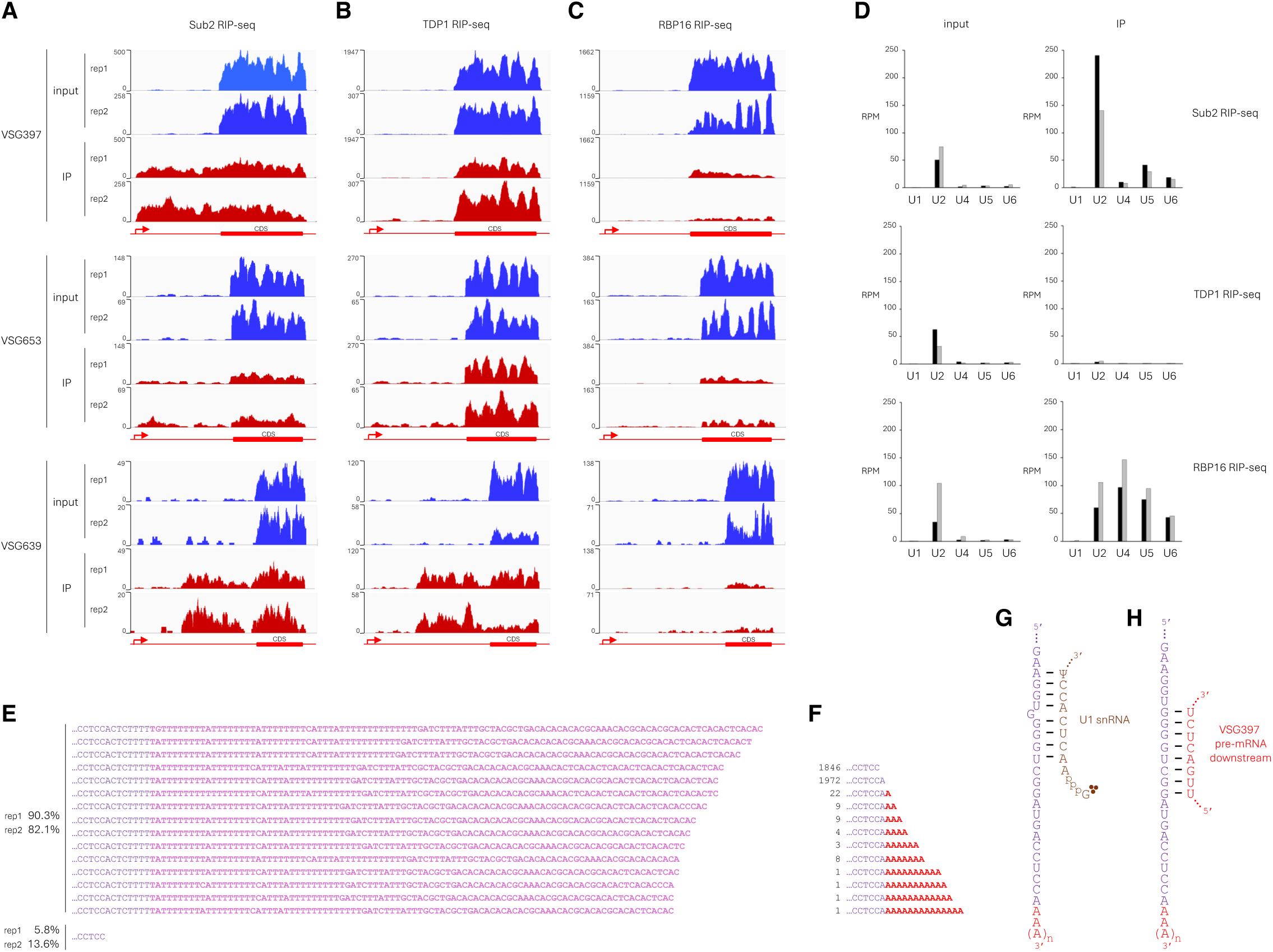
Sub2 associates with VSG397 pre-mRNA and precursor SL RNA. (A-C) RIP-seq analysis of Sub2 (A), TDP1 (B) and RBP16 (C). Reads are aligned to three metacyclic VSG expression sites exhibiting high (VSG397), intermediate (VSG653), and low (VSG639) frequencies of activation during differentiation to metacyclic forms. Data from two biological replicates (rep1 and rep2) are shown. IP, immunoprecipitate. (D) Reads mapping to spliceosomal U snRNAs in Sub2, TDP1, and RBP16 RIP-seq datasets. Black and grey bars represent the two replicates. RPM, reads per million. (E) Distribution of SL RNA 3′ ends detected in the two Sub2 RIP-seq replicates. The 15 longest genome-encoded 3′ extensions are highlighted in magenta. (F) Distribution of SL RNA 3′ ends in the U1C iCLIP dataset. Non-templated adenosine residues are shown in red. (G) Predicted base-pairing interaction between the 3′ end of SL RNA and the 5′ end of *T. brucei* U1 snRNA. (H) Predicted base-pairing interaction between the 3′ end of SL RNA and the VSG397 pre-mRNA downstream region.

The strong association of Sub2 with VSG pre-mRNA prompted us to examine reads mapping to the second substrate for SL *trans*-splicing, the SL RNA. Surprisingly, the SL RNA species associated with Sub2 were dominated by precursors carrying long genome-encoded 3′ extensions (Figure 3E). Reads containing such extensions accounted for 86.2 ± 4.1% (n=2) of reads spanning SL RNA positions 125–140 in the Sub2 immunoprecipitates, with the longest tails extending 95-110 nt beyond the known 3′ end of the 148-nt Sm proteins-assembled SL RNA (Figure 3E). These 3′ extensions are the result of continued transcription past the mature SL RNA 3′ end and consist of very U-rich first half and AC-dinucleotide dominated second half. In contrast to the 100% identity observed for the first 148-nt of the multiple SL RNA genes in *T. brucei*, the sequences past position +149 (relative to the transcription start site) are variable (Figure 3E). The combined length of these precursors (SL RNA plus tail) closely matches that of an SL RNA species previously detected following UV irradiation of *T. brucei* cells,^66^ which was originally presumed to represent a branched UV cross-linking adduct. Its identity likely remained unresolved because of sequence heterogeneity among SL RNA genes in the tail region (Figure 3E). The enrichment of tailed SL RNA precursors in Sub2 immunoprecipitates, together with the near absence of mature Sm-assembled SL RNA 3′ ends, indicates that the 3′-extended SL RNA precursors are bona fide substrates for pre-mRNA processing in *T. brucei*. The second most abundant SL RNA species in the Sub2 immunoprecipitates terminated at nucleotide C140 and represented 9.7 ± 3.9% (n=2) of all detected SL RNA 3′ ends. Because Sub2 associates with both substrates and products of pre-mRNA processing, the C140-terminated species may represent an end product of trypanosome mRNA maturation. Together, the identified 3′-tailed pre-SL RNA and C140-terminated SL RNA accounted for 95.9 ± 0.2% (n=2) of all Sub2-associated SL RNA detected by RIP-seq.

We next compared our findings with previous analyses of the interaction between SL RNA and the spliceosomal U1 snRNP.^18^ Examination of the most robust available U1C iCLIP dataset revealed numerous reads terminating at different positions near the 3′ end of SL RNA (Figure S5). Because the majority of U1C iCLIP reads terminated at the trinucleotide CCA (SL RNA positions 139-141), we focused our analysis on the central position, C140. In addition to reads terminating at C140 and possibly A141, we identified reads containing additional adenosine residues downstream of A141 (Figure 3F), indicating that U1 snRNP also associates with SL RNA molecules polyadenylated at the CCA trinucleotide. It remains unclear whether A141 represents the terminal nucleotide prior to poly(A)-tail addition or the first nucleotide of the synthesized poly(A) tail. These U1-associated polyadenylated SL RNA species were not detected in the Sub2 immunoprecipitates, which are enriched for pre-mRNA processing substrates and end products. We therefore infer that they likely represent short-lived intermediates in pre-mRNA/pre-SL RNA processing. CPSF73, the polyadenylation endonuclease,^67^ possesses a highly conserved active site among trypanosomes and exhibits a preference for cleavage at CA dinucleotides,^68,69^ making it an attractive candidate for catalyzing SL RNA 3′-end cleavage prior to polyadenylation. Because U1 snRNA uses its highly conserved 5′ end to recognize and base-pair with the 5′ splice site during *cis*-splicing,^70^ we examined the 3′ end of SL RNA and found that nucleotides 2-9 of *T. brucei* U1 have the potential to base-pair with a region of the SL RNA located 13 nt upstream of the CA cleavage site dinucleotide (Figure 3G). In addition, the VSG397 downstream region contains an 8-nt stretch complementary to nucleotides at the 3′ end of SL RNA located 10 nt upstream of the CA cleavage site (Figure 3H). This observation suggests that the VSG pre-mRNA region downstream of the conserved 16-mer and polyadenylation site may contribute directly to VSG mRNA maturation.

### Similarities between SL RNA and the VSG pre-mRNA downstream region to RNA elements required for RNA splicing and U-insertion/deletion pre-mRNA editing

The hypermethylated 5′ end of SL RNA in trypanosomatids, has the sequence 5′-AACUAA(C/A)-3′ (Figure S6), which closely resembles the branch-site sequence used during pre-mRNA splicing in eukaryotes with intron-poor genomes, such as *Saccharomyces cerevisiae* (5′-UACUA**A**C-3′) (Figure 4A) and *Giardia intestinalis* (5′-ACU[A/G)]**A**C-3′). Remarkably, a similar 5′-AACUAAA-3′ element is present in VSG pre-mRNAs immediately 5′ to the strong downstream polypyrimidine tract (Figure S7), the typical location of a pre-mRNA branch-site adenosine whose 2′-OH acts as the nucleophile in the first step of splicing. Downstream of the same polypyrimidine tract in VSG genes (Figure S7) lies an element with the potential to function as a branch-site recognition region (BSRR) (Figure 4B), a feature that is absent from trypanosomatid U2 snRNA.^11–15^ Additionally, the conserved 16-mer located at the extreme 3′ end of the VSG 3′UTR exhibits complementarity to the SL RNA exon and has the potential to form a secondary structure resembling both the bulge-interrupted duplex formed between guide RNA (gRNA) and pre-mRNA during mitochondrial U-insertion/deletion RNA editing^71^ (Figures 4B and 4C) and the base-pairing interactions involved in 5′ splice-site positioning during the first step of group II intron splicing^72^ (Figure 4D). Notably, RBP16, the top protein identified in the splicing enhancer pulldown assay, has been shown to facilitate annealing of the gRNA/pre-mRNA duplex and its subsequent rearrangement.^71^ The proposed interaction between the VSG 16-mer and the SL exon would bring into close proximity the SL RNA 5′ splice site, the cleavage site associated with polyadenylation, and the 5′-AACUAA-3′ element at the beginning of SL RNA (Figure 4B).

**Figure 4.**
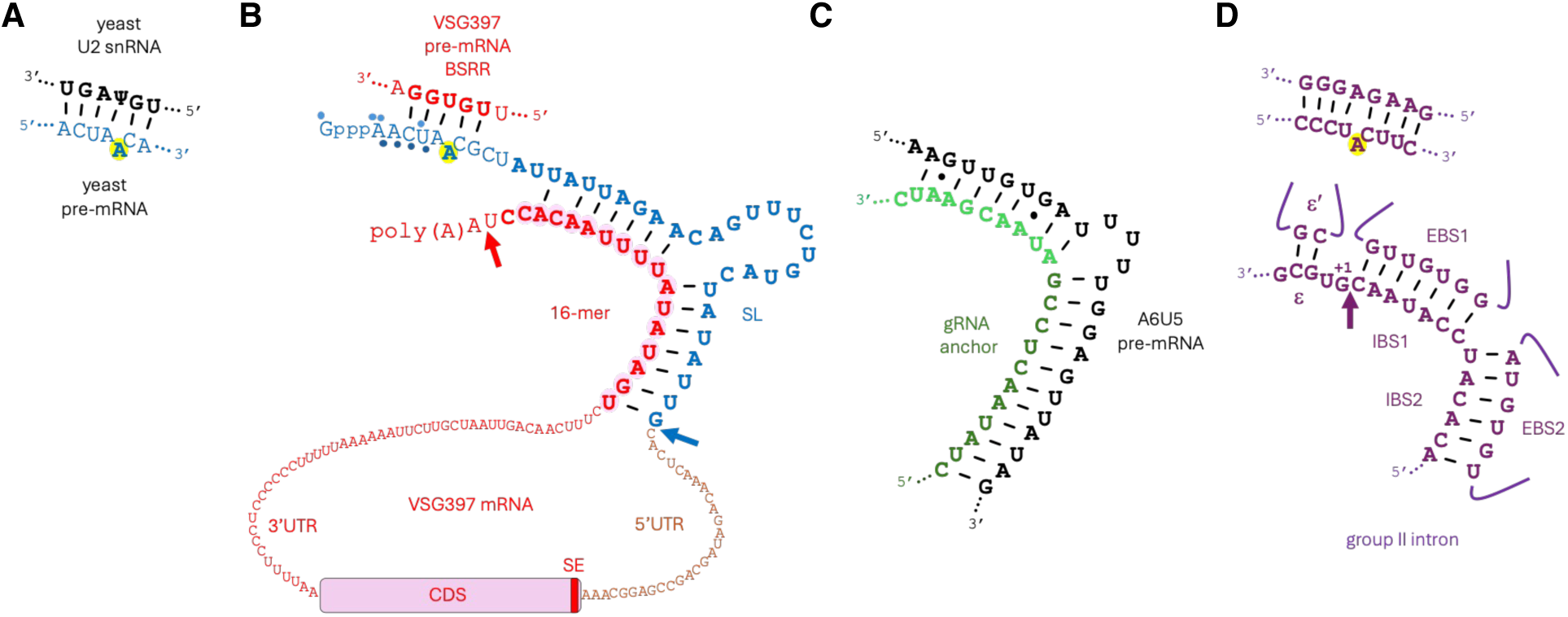
Sequence and structural similarities between proposed VSG pre-mRNA interactions and RNA-RNA contacts involved in splicing and mitochondrial pre-mRNA editing. (A) Base-pairing interaction between the *S. cerevisiae* U2 snRNA branch-site recognition region (BSRR) and the ACT1 pre-mRNA branch site. The branch-site adenosine is highlighted. (B) Predicted base-pairing interactions between the VSG397 pre-mRNA branch-site recognition region (BSRR) and the SL RNA 5′ end, and between the VSG 16-mer and the SL exon. The red arrow indicates the major polyadenylation site, and the blue arrow indicates the junction between the SL 5′ splice site and the VSG397 3′ splice site. (C) Interaction between guide RNA gA614NX and A6U5 pre-mRNA^71^ during U-deletion editing in the mitochondrion. (D) Base-pairing interactions involving the 5′ splice site and branch site during the first step of *Ll.LtrB* group II intron splicing.^72^ The arrow indicates the 5′ splice site.

In addition to the optimal 5′-AACUAA-3′ branch-site sequence, both SL RNA and the VSG pre-mRNA downstream region contain a U6-like catalytic triad, AGC or AAC (Figures 5A, 5B, S6 and S7). Although all known U6 RNAs contain an AGC at the spliceosomal catalytic center, substitution of the central G with A can support function in yeast^73^ and has no effect on the spliceosome activity in human cells,^74^ underscoring the requirement for a purine at this position. Similarities between *T. brucei* U6 snRNA and SL RNA also extend to the region between the U6 internal stem-loop (ISL) and the U6/U2 helix II (underlined in Figures 5A and 5B), which corresponds to the putative Sm-binding site in SL RNA.

**Figure 5.**
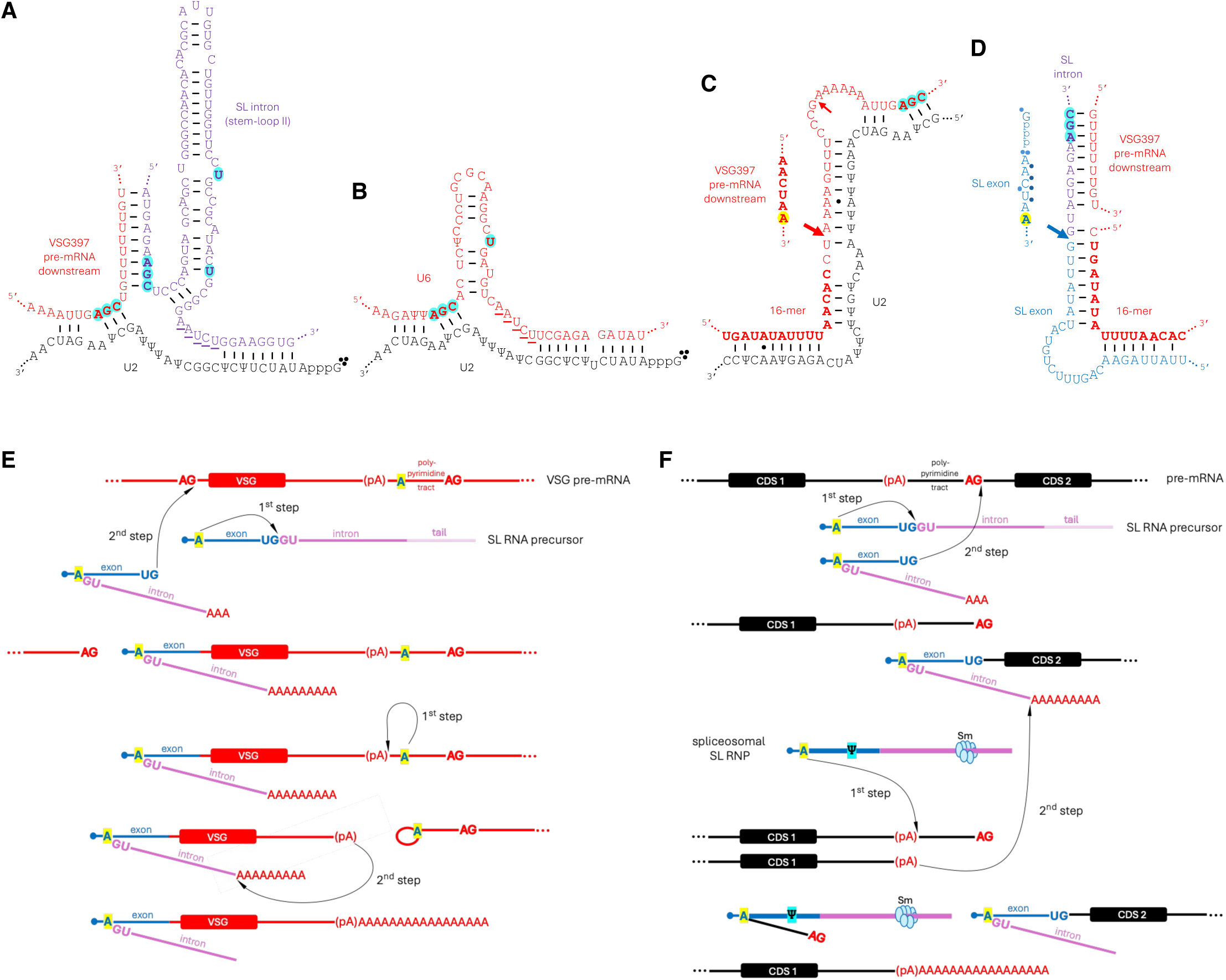
Model for VSG pre-mRNA processing. (A) Proposed catalytic core of the VSG pre-mRNA spliceosome. (B) Predicted base-pairing interactions between *T. brucei* U2 and U6 snRNAs in the *cis*-spliceosome. (C) Predicted base-pairing interactions that position the major VSG397 polyadenylation site (large red arrow) for cleavage by the VSG pre-mRNA spliceosome. The minor polyadenylation site is indicated by the small red arrow. (D) Predicted base-pairing interactions that position the SL RNA precursor 5′ splice site (blue arrow) for cleavage by the VSG pre-mRNA spliceosome. (E and F) Proposed mechanisms for SL and poly(A)-tail *trans*-splicing during processing of VSG pre-mRNA (E) and general trypanosome pre-mRNA (F). The SL RNA precursor undergoes CPSF73-catalyzed cleavage followed by polyadenylation by PAP2 at the beginning of the proposed processing cascade.

### Model for VSG and non-VSG pre-mRNA processing

Integrating our observations with published data on pre-mRNA processing in trypanosomatids, we propose an updated model for VSG mRNA maturation in *T. brucei*. Because the conserved 16-mer in the VSG 3′UTR has potential to engage in base-pairing interactions with both U2 snRNA (Figure 5C) and the SL exon (Figure 5D), we propose that it acts as a central organizing element in a network of spliceosomal RNA-RNA interactions that drive SL *trans*-splicing and spliceosome-catalyzed transfer of poly(A) tail from a polyadenylated SL RNA precursor. In this model, cleavage and polyadenylation of an SL RNA precursor carrying a long 3′ extension constitutes an essential early step in the coupled processes of SL addition and polyadenylation.

The model involves two successive rounds of *trans*-splicing (Figure 5E). During the first step of the first (SL) round, the 2′-OH of the branch-site adenosine within 5′-AACUA**A**-3′ sequence at the 5′ end of the substrate SL RNA attacks the 5′ splice site intramolecularly, generating a Y-structure intermediate. In the second step of SL *trans*-splicing, the exposed 3′-OH of the SL exon attacks the 3′ splice site in the VSG pre-mRNA, resulting in formation of the mature SL-containing 5′ end of the transcript. During the first step of the second [poly(A)] round of *trans*-splicing, the 2′-OH of the branch-site adenosine within the 5′-AACUA**A**-3′ element adjacent to the downstream VSG polypyrimidine tract attacks the polyadenylation site, generating a lariat intermediate. In the second step of poly(A) *trans*-splicing, the liberated 3′-OH at the VSG polyadenylation site attacks the poly(A) tail of the SL RNA substrate, thereby joining the VSG mRNA body to a poly(A) tail derived from the SL RNA precursor.

For the general mechanism of non-VSG pre-mRNA processing (Figure 5F), SL *trans*-splicing proceeds through a pathway analogous to that proposed for VSG mRNA maturation. Because non-VSG genes lack a conserved 5′-AACUAA-3′ branch-site element for poly(A) *trans*-splicing, we propose that the Sm-assembled, Ψ28-containing SL RNA functions as a bona fide spliceosomal RNA rather than as an SL donor. Its primary role is to provide the branch-site adenosine within the 5′-AACUA**A**-3′ sequence that serves as the nucleophile for attack on the pre-mRNA polyadenylation site. The second step of poly(A) *trans*-splicing is proposed to proceed similarly to that of VSG mRNA maturation. The involvement of two distinct SL RNA molecules, a substrate precursor that undergoes cleavage and polyadenylation, and a genuine Sm-containing spliceosomal SL RNP that contributes a second branch-site adenosine for poly(A) *trans*-splicing, provides an explanation for why mature trypanosome mRNAs contain the co-transcriptionally acquired hypermethylated cap, but evidently lack Ψ28 within the transferred SL sequence.

### Extensive complementarity between the VSG397 splicing enhancer and the VSG397 pre-mRNA downstream region

Trypanosomes (class Kinetoplastea) belong to the phylum Euglenozoa, which also includes the classes Euglenida and Diplonemea. In addition to conventional introns, the genomes of both diplonemids and euglenids contain nonconventional introns, characterized by noncanonical boundary sequences and extensive base-pairing interactions that bring the 5′ and 3′ splice site into close proximity.^75,76^ This observation prompted us to investigate the possibility for additional interactions between the substrates involved in SL and poly(A)-tail *trans*-splicing of VSG397 pre-mRNA. We focused on the regions surrounding the putative branch-site recognition region (BSRR) in VSG397 pre-mRNA, as they are located adjacent to the proposed spliceosomal active site. We found that the region downstream of the BSRR, which contains degenerate telomeric (subtelomeric) repeats, has the potential to base-pair with the 3′ end of the SL RNA (Figure 6A). The highly U-rich polypyrimidine tract immediately upstream of the BSRR also has the potential to interact with the poly(A) tail of the substrate SL RNA, although the precise base-pairing register is difficult to predict (Figure 6A). Interestingly, the same region encompassing the polypyrimidine tract, BSRR, and subtelomeric repeats has the potential to base-pair both with the sequence downstream of the VSG397 3′ splice site through the polypyrimidine tract and with the splicing enhancer located at the beginning of the VSG397 CDS through the subtelomeric repeats (Figure 6B). Although the subtelomeric repeats differ in sequence among VSG genes, they retain the capacity to form analogous interactions with the corresponding VSG coding sequence, which themselves are not homologous (Figure 6C). Analysis of the two most highly expressed sequences in all three replicates of the *in vivo* mutagenized VSG397 splicing enhancer screen revealed an increased potential for base-pairing in the immediate vicinity of the BSRR (Figure 6D). In contrast, a sequence that was the most abundant before selection (day 0) but absent following induction of metacyclic differentiation (day 6) exhibited a reduced capacity for interaction near the BSRR (Figure 6E). Consistent with these observations, the sequence of the 3′ half of the splicing enhancer conformed to a higher degree to the wild type VSG CDS (Figure 1E), supporting the idea that interactions between the splicing enhancer and the subtelomeric repeats adjacent to the BSRR contribute to efficient VSG397 pre-mRNA processing.

**Figure 6.**
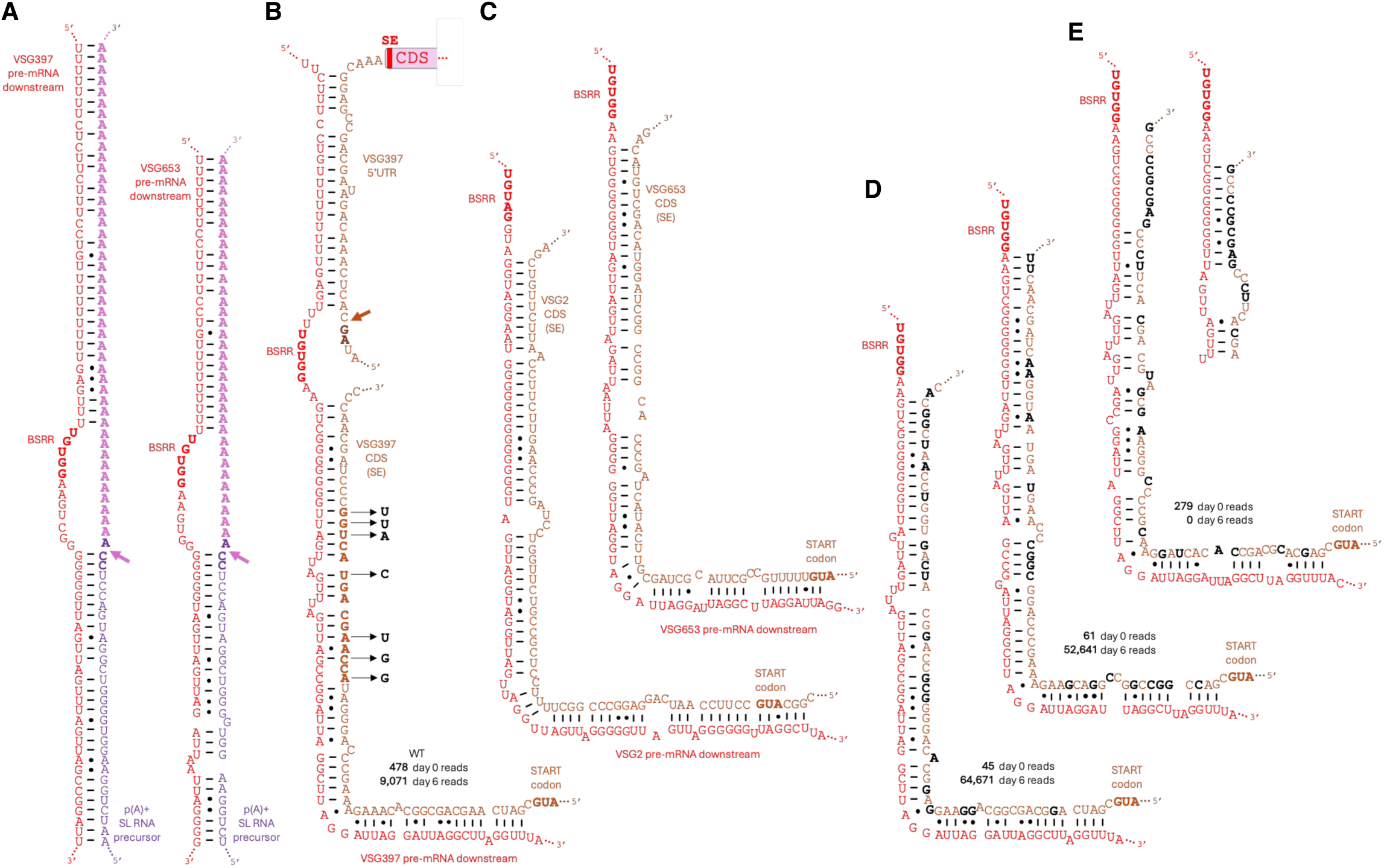
Potential VSG pre-mRNA base-pairing interactions involved in substrate positioning during the second step of poly(A)-tail and SL *trans*-splicing. (A) Predicted base-pairing interactions between the downstream regions of VSG397 and VSG653 pre-mRNAs and the polyadenylated 3′ end of precursor SL RNA. Non-canonical A•G, A•A, G•G, and U•U base pairs are indicated. A•C and U•C base pairing is possible, but not denoted. The purple arrow indicates the predicted 3′ splice site for poly(A)-tail *trans*-splicing. (B) Predicted base-pairing interactions between the downstream region of VSG397 pre-mRNA, the SL addition site (brown arrow indicates the 3′ splice site), and the splicing enhancer (SE) located at the beginning of the VSG397 CDS. The 15-nt RNA segment used in the protein pulldown assay is shown in bold, and the mutated nucleotides are indicated. The numbers of wild-type reads recovered on days 0 and 6 of induced metacyclic differentiation in the mutagenized SE library screen are shown. (C) Predicted base-pairing interactions between the downstream regions of VSG2 and VSG653 pre-mRNAs and their corresponding splicing enhancers. (D) Predicted interactions between the two most abundant mutant SE sequences identified in the mutagenized library screen and the VSG397 pre-mRNA downstream region. Mutated residues are shown in black. (E) Predicted interactions between the VSG397 pre-mRNA downstream region and a mutant SE sequence that is absent on day 6 of induced metacyclic differentiation but abundant in the mutagenized SE library on day 0. Two alternative base-pairing interactions in the vicinity of the branch-site recognition region (BSRR) are shown. Mutated residues are shown in black.

### CSD1 interacts with the VSG397 pre-mRNA subtelomeric repeats and branched SL RNA

Previous analysis of mRNA binding by CSD1^40^ concluded that the protein does not interact with mature VSG mRNA, consistent with its absence from purified VSG mRNPs.^34^ As expression of CSD1 (or CSD2) is the most rapid known trigger of metacyclic VSG expression,^40^ we hypothesized that the protein may participate in nuclear RNA metabolism and therefore examined published CSD1 seCLIP data^40^ for evidence of interactions with VSG397 pre-mRNA. We confirmed the near absence of seCLIP reads mapping to the VSG397 5′UTR, CDS or 3′UTR. Although no reads were detected upstream of the VSG397 3′ splice site for SL addition, reads mapping to the subtelomeric repeats downstream of the putative VSG397 BSRR were readily observed (Figure 7A), suggesting a role for CSD1 in VSG397 pre-mRNA processing. Analysis of the CSD1 seCLIP dataset with respect to small non-coding RNAs further revealed associations with the SL RNA (both exon and intron sequences), U2 snRNA, and vtRNA, a small RNA implicated in SL *trans*-spliced mRNA production in *T. brucei*^77^ (Figure 7B). Inspection of the CSD1-bound SL exon reads revealed that they most frequently terminated at nucleotide A5, one nucleotide upstream of the proposed branch-site adenosine A6 in the highly conserved 5′-AACUA**A**-3′ (Figure 7C). In addition, a conspicuous number of reads contained an A6-to-C substitution (Figure 7D), a known signature of reverse transcriptase traversing a branched nucleotide in RNA.^78^ This substitution was not observed at any other adenosine residue in the first half of SL RNA (Figure 7D). Analysis of CSD2 seCLIP dataset yielded highly similar results (not shown). Together, these observations support the prediction that A6 in SL RNA functions as a branch-site adenosine and are consistent with our proposed model of *T. brucei* pre-mRNA processing (Figure 5). Moreover, these findings suggest that RT primer-extension assays of the SL RNA exon do not exclusively report on the methylation status of the SL RNA 5′ end, as has often been assumed.^79^ Consequently, previous studies that interpreted reduced RT termination at A5 solely as evidence of altered SL cap hypermethylation may warrant re-interpretation in light of a potential contribution from branch-site-dependent RT stops.

**Figure 7.**
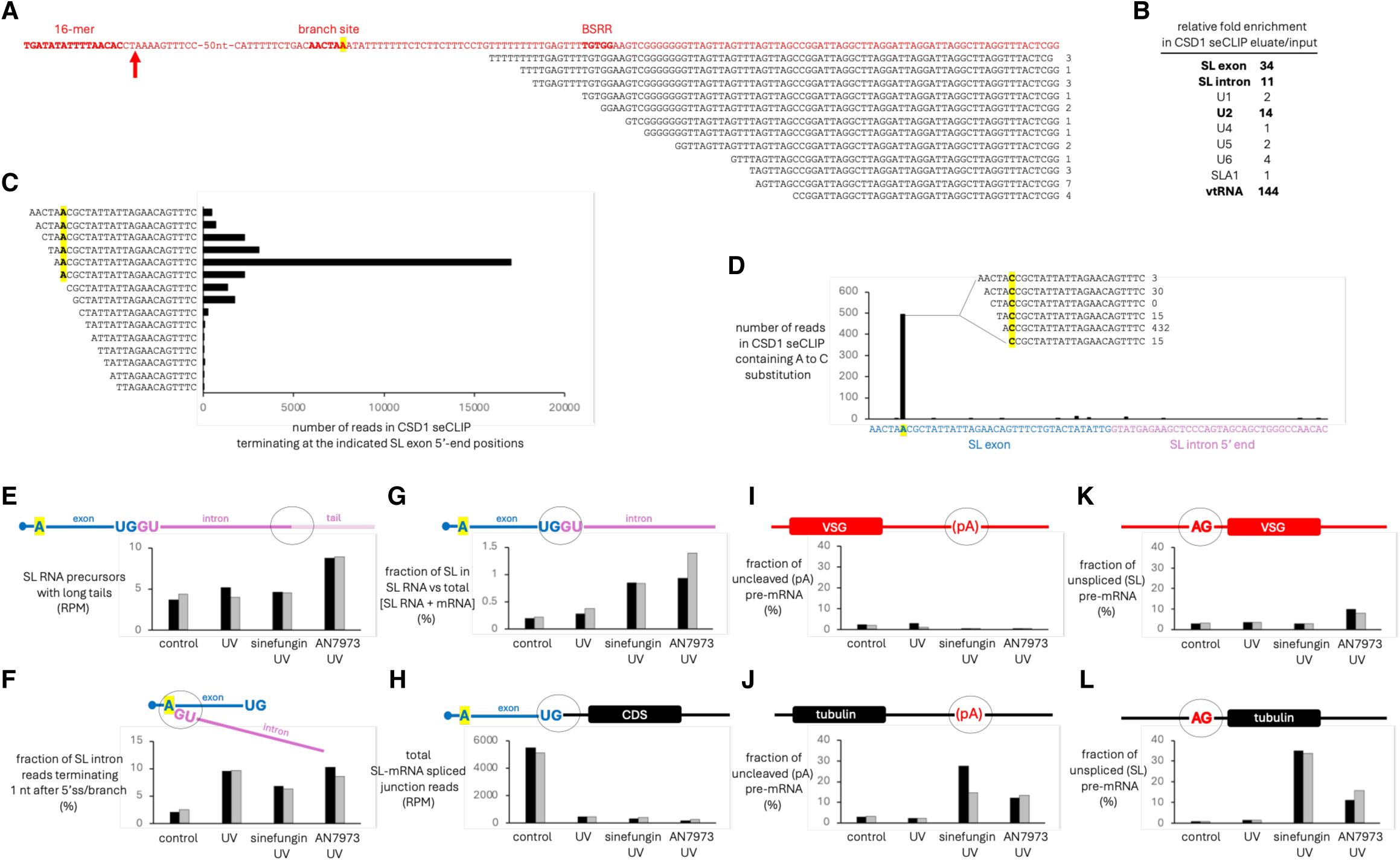
CSD1 associations with splicing intermediates and limited effects of pre-mRNA processing inhibitors on VSG mRNA maturation. (A) CSD1 seCLIP reads mapping to the VSG pre-mRNA downstream region. Numbers indicate read counts. (B) Enrichment of spliceosomal snRNAs in the CSD1 seCLIP eluate relative to input. 5S rRNA was used for normalization. (C) 5′ ends of CSD1 seCLIP reads mapping to the SL exon. The predicted branch-site adenosine is highlighted. (D) Distribution of CSD1 seCLIP reads mapping to the SL exon and containing A-to-C substitutions. (E)-(L) Effects of UV irradiation, sinefungin (methylation inhibitor), and AN7973 (CPSF73 inhibitor) on the abundance of pre-mRNA processing substrates, intermediates, and products in *T. brucei*. Black and gray bars represent two biological replicates.

### Limited effect of the methylation inhibitor sinefungin and the CPSF73 inhibitor AN7973 on VSG pre-mRNA processing

The proposed models for VSG and non-VSG mRNA maturation differ primarily in the identity of the RNA that contributes the branch-site adenosine for poly(A)-tail *trans*-splicing. In the VSG model, an adenosine upstream of the polypyrimidine tract adjacent to the subtelomeric repeats in VSG pre-mRNA serves as the nucleophile for the first transesterification reaction that cleaves the polyadenylation site, whereas in the general model the branch-site adenosine is provided by the 5′ end of the spliceosomal SL RNA (Figure 5E,F). There is currently no information regarding the methylation status of the 5′-AACUA**A**-3′ sequence in VSG pre-mRNA, and this region of the transcript is only rarely detected in RNA-seq data, precluding direct analysis. Nevertheless, we consider it unlikely that the same methylations found on SL RNA are present in VSG pre-mRNA, because these nucleotides are not adjacent to a m^7^G(5’)ppp(5’)N cap structure. Consequently, a nonmethylated 5′-AACUA**A**-3′ should remain functional in the presence of sinefungin, a methylation inhibitor known to inhibit *T. brucei* pre-mRNA processing.^79^ To test this prediction, we treated cells induced to differentiate into metacyclic forms and express VSG genes with either sinefungin or the CPSF73-specific inhibitor AN7973,^7^ followed by UV irradiation to increase the likelihood of detecting splicing substrates and intermediates. UV-induced cross-linking is expected to inhibit spliceosomal rearrangements and thereby stabilize otherwise transient processing intermediates. Total RNA and poly(A)+ RNA were subsequently analyzed by RNA-seq.

AN7973 treatment caused a two-fold increase in the abundance of SL RNA precursors carrying a long genome-encoded tail, whereas sinefungin or UV irradiation alone had little effect (Figure 7E). These observations support the prediction that CPSF73 is responsible for cleavage of pre-SL RNA prior to polyadenylation. Sinefungin reduced the proportion of reads terminating one nucleotide upstream of the 5′ splice site, the predominant reverse-transcription stop in these libraries. This population includes reads corresponding to Y-structure intermediates, as reverse transcriptase preferentially terminates one nucleotide after a branched residue. The reduction was observed relative to both UV-only and AN7973/UV-treated samples (Figure 7F). These results are consistent with inhibition of SL intron branching resulting from impaired methylation of the SL RNA 5′ end, which is generally required for *trans*-splicing.^79^ The proportion of SL exon sequences present in SL RNA relative to total SL-containing RNA (SL RNA plus mRNA) increased similarly following treatment with either sinefungin or AN7973 (Figure 7G), indicating that CPSF73 activity is required for substrate SL RNA participation in SL *trans*-splicing. UV treatment alone resulted in a marked reduction in total SL-*trans*-spliced mRNA (Figure 7H), consistent with observations that UV cross-linking substantially slows RNA splicing.

Comparison of the effects of sinefungin and AN7973 on 3′-end cleavage of VSG and tubulin pre-mRNAs revealed that neither treatment inhibited 3′-end processing of VSG pre-mRNA (Figure 7I). In contrast, uncleaved tubulin pre-mRNA accumulated to similar levels in both sinefungin- and AN7973-treated cells (Figure 7J), indicating that tubulin pre-mRNA 3′-end cleavage requires both CPSF73 activity and methylation of the SL RNA 5′ end. Likewise, sinefungin had no detectable effect on 5′-end processing of VSG397 pre-mRNA (Figure 7K), whereas inhibition of tubulin SL *trans*-splicing was readily apparent, resulting in accumulation of unspliced tubulin pre-mRNA to 34.3 ± 0.7% (n=2) (Figure 7L). In contrast, AN7973 negatively affected SL *trans*-splicing of both VSG397 and tubulin pre-mRNAs (Figure 7K,L).

Together, these findings are consistent with the proposed models for *T. brucei* pre-mRNA processing (Figure 5E,F) and suggest that CPSF73-mediated cleavage of pre-SL RNA is a prerequisite for SL *trans*-splicing. Furthermore, the markedly different sensitivities of VSG and tubulin pre-mRNA processing to sinefungin support the existence of two distinct processing mechanisms. In the general model, both SL and poly(A)-tail *trans*-splicing require methylated SL RNA 5′ ends to provide branch-site adenosines (Figure 5F). In contrast, the spliceosomal machinery responsible for VSG pre-mRNA processing appears capable of functioning in the absence of methylation because it utilizes a branch-site adenosine within the pre-mRNA itself during 3′-end processing (Figure 5E).

## DISCUSSION

The model for VSG pre-mRNA processing presented here involves the participation of the precursor mRNA itself as an integral component of a specialized spliceosome, which we term the VSG pre-mRNA spliceosome. In this model, the VSG downstream region contributes a catalytic A(G/A)C triad, whereas the conserved 16-mer at the end of the 3′UTR acts as a central organizing element within a dynamic spliceosomal base-pairing interactions network. The 16-mer and the VSG pre-mRNA downstream region are proposed to direct both SL and poly(A)-tail *trans*-splicing through interactions with the SL RNA substrate and U2 snRNA. In addition, the downstream region has the potential, analogous to U1 snRNA, to base-pair with the 3′ end of the substrate SL RNA and thereby direct its cleavage and polyadenylation. Thus, the VSG pre-mRNA spliceosome may represent a streamlined and highly efficient processing apparatus that does not require the participation of U1, U6 or the spliceosomal SL RNA. In our model, both the SL sequence and the poly(A) tail are transferred from a polyadenylated SL RNA substrate through two successive rounds of *trans*-splicing, each involving the two transesterification reactions characteristic of canonical pre-mRNA splicing. This mechanism provides a framework for reconciling several uncertainties associated with the classical model of SL *trans*-splicing in *T. brucei* and offers an explanation for the tight coupling of 5′- and 3′-end formation in trypanosomatid mRNAs. Furthermore, our findings suggest that evolution may have produced distinct molecular architectures capable of carrying out the same two transesterification reactions, preserving the fundamental chemistry of RNA splicing while allowing considerable variation in the RNA components responsible for catalysis and substrate positioning.

The involvement of two SL RNA molecules in the maturation of non-VSG transcripts—one functioning as the substrate and the other as a bona fide spliceosomal snRNA—suggests that the Sm-assembled SL RNP plays an important role in the process, potentially by positioning RNA substrates for the second steps of both SL and poly(A)-tail *trans*-splicing (Figure S8), in addition to providing a branch site adenosine during poly(A)-tail *trans*-splicing. Such a role would be analogous to the proposed function of the subtelomeric repeats in VSG pre-mRNA. With regard to activation of the 5′ splice site in the substrate SL RNA, two small RNAs with established links to SL RNA metabolism and *trans*-splicing in *T. brucei* emerge as attractive candidates: SLA1 RNA (spliced leader-associated RNA 1)^80^ and vtRNA (vault RNA)^77^ (Figure S9). Like mammalian telomerase RNA,^81^ SLA1 is stabilized by box H/ACA RNP proteins. Although SLA1 is able to guide Ψ28 formation within the spliceosomal SL RNA exon,^23^ its psoralen cross-linking pattern to SL RNA^80^ suggests an alternative mode of interaction (Figure S9), raising the possibility that, similar to the box H/ACA RNP associated with human telomerase RNA, pseudouridylation may not be its primary function. The involvement of vtRNA in SL *trans*-splicing has been demonstrated previously.^77^ Both SLA1 RNA and vtRNA contain potential BSRRs for the SL RNA 5′-AACUA**A**-3′ element and exhibit substantial complementarity to the SL RNA 5′ splice site (Figure S9). Together, these observations raise the intriguing possibility that trypanosomes possess multiple classes of *trans*-spliceosomes, analogous to the coexistence of the major (U2-type) and minor (U12-type) spliceosomes in other eukaryotes.^82^ Such complexes could include vtRNA-type, SLA1-type, and VSG pre-mRNA-type *trans*-spliceosomes, in addition to the conventional *cis*-spliceosome composed of U1, U2, U4, U5, and U6 snRNPs.

Our analyses suggest that the primary role of U1 snRNA in trypanosome *trans*-splicing is interaction with the 3′ end of the substrate SL RNA to direct cleavage by CPSF73 and subsequent polyadenylation. Although a stimulatory role for U1 in small RNA polyadenylation has not been described previously, extensive functional connections between U1 snRNA and the cleavage/polyadenylation machinery are well established. The best-characterized example is the ability of mammalian U1 snRNA to base-pair close to cryptic polyadenylation signals and thereby suppress premature cleavage and polyadenylation^83^ or regulate polyadenylation-site choice^84^ through interactions with the cleavage/polyadenylation machinery.^85^ Our analyses indicate that *T. brucei* U1 snRNP promotes cleavage and polyadenylation of SL RNA substrates, a role reminiscent of U7 snRNP,^86^ which directs CPSF73-mediated 3′-end cleavage of non-polyadenylated histone mRNAs in metazoans. Interestingly, *T. brucei* U1 snRNA is unusually short (73 nt), a length comparable to that of U7 snRNA.

The identification of two proteins involved in mitochondrial U-insertion/deletion RNA editing, RBP16 and DRBD18, as factors associated with the VSG397 splicing enhancer and, in the case of RBP16, the splicing machinery itself, further highlights previously recognized nuclear functions of auxiliary mitochondrial RNA-editing factors.^48–50^ These proteins appear particularly well suited to facilitate dynamic RNA-RNA interactions involving the formation and subsequent remodeling of imperfect helices containing small bulges. A notable example is the proposed interaction between the VSG splicing enhancer and the downstream subtelomeric repeats within VSG pre-mRNA. Together, these findings support the existence of functional (and potentially evolutionary) connections between mitochondrial RNA editing and nuclear pre-mRNA *trans*-splicing. They also suggest an unexpected role for mitochondrial RNA editing in maintaining expression and activity of RNA-binding proteins required for VSG mRNA maturation, even during life-cycle stages in the insect vector in which VSG genes are not expressed.

More broadly, the model proposed here highlights the deep evolutionary relationship between RNA splicing and 3′-end formation (Figure S10). Additional evidence for this connection comes from observations that spliceosome-mediated cleavage generates the 3′ end of telomerase RNA in fission yeasts and filamentous fungi through a reaction mechanistically equivalent to the first transesterification step of pre-mRNA splicing.^87–89^ Furthermore, Integrator complex subunit 11 (INTS11), the endonuclease responsible for formation of snRNA 3′ ends in higher eukaryotes, is a paralog of CPSF73.^90,91^ Their shared evolutionary origin is reflected in our model, in which CPSF73 mediates cleavage of the SL RNA precursor prior to polyadenylation. Although computational analyses have not yet resolved questions surrounding the earliest evolution of eukaryotic pre-mRNA splicing, the findings presented here from an extant early-diverging eukaryote suggest that *trans*-splicing and spliceosome-catalyzed poly(A)-tail addition represent an ancient strategy for eukaryotic pre-mRNA processing in a genome where *cis*-introns are almost absent.

## Supporting information

Instructions for original code

Original code

## ACKNOWLEDGMENTS

We thank Joan A. Steitz, Susan J. Baserga, T. Nicolai Siegel, and members of their laboratories for valuable feedback. We are grateful to J.A. Steitz, Kazimierz T. Tycowski and T.N. Siegel for their insightful comments on the manuscript. We thank Mark J. Byers for bioinformatics support and Katie Henke for advice on sample preparation for LC-MS/MS and for data processing. We acknowledge the Yale Center for Genome Analysis for Illumina sequencing, supported by the National Institute of General Medical Sciences (NIGMS) award S10OD030363-01A1, and the Yale Keck Mass Spectrometry & Proteomics Resource for LC-MS/MS analyses. The Yale Keck facility is supported in part by the Yale School of Medicine and by NIGMS award S10OD018034. This work was funded by the National Institute of Allergy and Infectious Diseases of the National Institutes of Health grant R01AI165480 (to C.T.).

## DATA AND CODE AVAILABILITY

- Illumina sequencing data have been deposited at the sequence read archive (SRA) and are publicly available with BioProject accession numbers PRJNA1499860, PRJNA1503010, and PRJNA1505653.
- Mass spectrometry data have been deposited to the ProteomeXchange Consortium via the PRIDE repository and are publicly available with identifier PXD082003.
- Original code is included as Supplemental Information.

## METHODS

### *T. brucei* cell lines and culture

Procyclic *T. brucei rhodesiense* YTat 1.1 cells were used for integration of reporter constructs into bloodstream expression sites. *T. brucei* Lister 427 29-13 cells inducibly expressing RBP6^39^ were used for VSG397 CDS deletion analyses, the doped mutagenesis screen of the VSG397 splicing enhancer, GFP-fusion localization assays, RIP-seq analyses, and RNA-seq following sinefungin, AN7973, and UV-irradiation treatments. *T. brucei* Lister 427 29-13 cells inducibly expressing CSD1^40^ were used to prepare extract for the splicing enhancer pulldown assay. Cells were maintained at 28°C in 5% CO2 in Cunningham’s medium supplemented with 10% Tet-system approved heat-inactivated fetal bovine serum (Clontech), 2 mM L-glutamine, 100 units/mL penicillin, 100 μg/mL streptomycin, and 50 μg/mL gentamicin. Selection drugs were used at the following concentrations: 15 μg/mL G418, 50 μg/mL hygromycin B, 2.5 μg/mL phleomycin, 10 μg/mL blasticidin, and 1 μg/mL puromycin. For each transfection, 1 × 10^8^ cells were collected by centrifugation, washed with Cytomix buffer (25 mM HEPES, 20 mM KCl, 10 mM K_2_HPO_4_, 2 mM EGTA, 5 mM MgCl_2_, 0.15 mM CaCl_2_, pH 7.6), resuspended in 0.5 mL Cytomix buffer and mixed with either 20 μg of linearized plasmid DNA or 1-2 μg of PCR product. Cells were electroporated using a Gene Pulser Xcell (Bio-Rad) with two pulses of 1,600 V and a time constant of 0.6 ms. Following electroporation, cells were transferred to 10 mL of fresh medium, and selection drugs were added 16 h later. Expression of CSD1 or RBP6 was induced with 10 μg/mL doxycycline.

### DNA constructs

All DNA constructs generated and used in this study are shown schematically in Figure S1.

### RNA isolation and northern blotting

Total RNA was isolated using TRIzol reagent (Invitrogen) according to the manufacturer’s instructions. RNA was separated on 1.2% agarose gels containing 6.3% formaldehyde in MOPS buffer (40 mM MOPS, 20 mM sodium acetate, 2 mM EDTA, pH 7.0). RNA was transferred to Amersham Hybond-N nylon membranes (GE Healthcare) by capillary blotting in 10× SSC (1.5 M sodium chloride, 0.15 M sodium citrate) for 24 h and UV crosslinked using an XL-1500 UV crosslinker (Spectronics Corporation). Ribosomal RNA was visualized by staining with methylene blue (0.03% in 0.3M sodium acetate, pH 5.4), followed by destaining with water. Membrane were prehybridized for 1h in a solution containing 5× SET (0.75 M sodium chloride, 5 mM EDTA, 0.15 M Tris-HCl, pH 7.4), 10× Denhardt’s solution, 1% SDS and 100 μg/mL yeast RNA. Hybridization with radiolabeled probes was carried out for 16 h in the same solution at 65°C. Membranes were washed twice for 20 min in wash buffer (2× SSC, 0.1% SDS) at 65°C. Hybridization signals were detected using BAS-IP MS 2025 storage phosphor screens (Fujifilm), scanned on a Typhoon FLA7000 (GE Healthcare), and analyzed with ImageQuantTL software (GE Healthcare).

### Doped mutagenesis splicing enhancer library screen

The mutagenized splicing enhancer library was generated by PCR using an oligonucleotide containing the doped sequence (Integrated DNA Technologies) as a forward primer (100 nM) in eight 100-μL reactions with Phusion DNA polymerase (New England Biolabs) for 23 cycles. The PCR products were purified and digested with NheI and NotI (New England Biolabs). Five micrograms of the digested insert were ligated to 2 μg of plasmid vector (Figure S1C), digested with the same restriction enzymes, in a 220 μL reaction containing T4 DNA ligase (Roche) for 14 h at 16°C. The ligation mixture was electroporated into MegaX DH10B *E. coli* cells (Invitrogen). Library quality was assessed by sequencing plasmid DNA isolated from 19 individual bacterial colonies. Approximately 1.2 × 10^7^ colonies were pooled, and plasmid DNA was isolated using the QIAGEN Plasmid Maxi kit. The mutagenized splicing enhancer plasmid library was digested with XhoI and NotI (New England Biolabs), and the DNA fragment used for transfection into *T. brucei* (Figure S1C) was purified from a 1% agarose gel. Approximately 3.6 × 10^9^ procyclic *T. brucei* Lister 427 29-13 cells inducibly expressing RBP6^39^ were transfected by electroporation with the purified library using 20 μg DNA per transfection in 36 independent electroporations (1 × 10^8^ cells per transfection). Cells were allowed to recover for 24 h before selection with 10 μg/mL blasticidin for 12 days.

Genomic DNA was isolated from 2 × 10^8^ trypanosomes cultured without RBP6 induction using the DNeasy Blood & Tissue Kit (QIAGEN). A parallel culture containing 2 × 10^8^ cells was induced with 10 μg/mL doxycycline for 6 days to express RBP6 and differentiate into metacyclic forms, after which total RNA was isolated using TRIzol reagent (Invitrogen). cDNA was synthesized using random hexadeoxynucleotide primers (Promega) and SuperScript III reverse transcriptase (Invitrogen) according to the manufacturer’s instructions. The splicing enhancer region was amplified from the initial plasmid library and genomic DNA from uninduced cells (day 0) using Phusion DNA polymerase (New England Biolabs), forward primer 5′-GGTGGCGTAAGTATTGCCTAATG-3′, and VSG397 reverse primer 5′-ATTGCGGAATTTATTGTGCTGCG-3′. Amplification of expressed VSG397 splicing enhancer variants from day 6 of RBP6 induction used cDNA as the template, the SL oligonucleotide 5′-TAGAACAGTTTCTGTACTATATTG-3′ as the forward primer, and the same VSG397 reverse primer. Illumina sequencing libraries were prepared from purified PCR products and sequenced at the Yale Center for Genome Analysis. Sequence logos were generated using WebLogo 3 (https://weblogo.berkeley.edu/).

### SE pulldown assay

One liter of a culture of procyclic-form *T. brucei* inducibly expressing CSD1^40^ (6 × 10^6^ cells/mL) was treated with 10 μg/mL doxycycline for 36 h. Cells were harvested by centrifugation and washed three times by resuspension in ice-cold PBSG (phosphate-buffered saline containing 25 mM glucose). Two cell pellets (approximately 0.2 mL each) were lysed in 1 mL of ice-cold buffer containing PBS, 1% NP-40, 5% glycerol, 2 mM DTT, 25 mM glucose, and Complete Mini EDTA-free protease inhibitor cocktail (Roche; one tablet per 5 mL). Following centrifugation at 13,200 rpm for 15 min at 4°C in an Eppendorf 5424 R microcentrifuge, the clarified lysates were transferred to fresh tubes, aliquoted (0.5 mL), snap-frozen on dry ice, and stored at −80°C.

For each pulldown replicate, a 0.5 mL aliquot of extract was thawed and mixed with an equal volume of ice-cold PBS. The extract was precleared by incubation with 0.2 mL of Dynabeads MyOne Streptavidin C1 slurry (10 mg/mL; Invitrogen) for 1 h at 4°C with gentle rotation. During this incubation, 2 μg of either the wild-type or mutant 3′-biotinylated 15-nt RNA bait (Dharmacon) were separately coupled to 50 μL of Dynabeads MyOne Streptavidin C1 slurry in 0.5 mL PBS for 1 h at 4°C with gentle rotation. Following magnetic separation of the preclearing beads, the extract was supplemented with 50 μL of 200 mM ribonucleoside vanadyl complex (New England Biolabs). The RNA-coupled beads were washed twice with 1 mL of ice-cold PBS, incubated with 0.5 mL of the supplemented extract for 1.5 h at 4°C with gentle rotation, and then washed five times with 1 mL of PBS containing 0.2% NP-40 followed by five washes with 1 mL of PBS alone. The beads were finally resuspended in 50 μL PBS and submitted to the Yale Keck Mass Spectrometry & Proteomics Resource for LC-MS/MS analysis using standard protocols for protein elution and tryptic digestion.

### Proteomics data analysis

Protein identification was performed in Proteome Discoverer (Thermo Fisher Scientific, version 3.0) using standard processing and consensus workflows. Briefly, precursor masses were recalibrated before data were searched against the UniProt *Trypanosoma brucei brucei* database (9,426 entries; downloaded November, 2024) and a database of common contaminant proteins. The CHIMERYS^1^ 1 search engine was employed using the INFERYS^2^ 2.1 prediction model. Trypsin was selected as the enzyme, up to two missed cleavages were allowed, and the fragment ion mass tolerance was set to 0.02 Da. Peptides with lengths between 7-30 amino acids and charge states +1-6 were allowed. Oxidation of methionine was set as a variable modification and carbamidomethylation of cysteine was set as a fixed modification. The Minora algorithm was used for feature detection and PSM-level false discovery rates (FDRs) were estimated using Percolator^3^. Scaffold DDA (Proteome Software Inc., version 6.6.0) was used to filter peptide and protein identifications. Peptide- and protein-level FDR thresholds were set to 1% and only proteins identified by at least two peptides were considered for downstream analysis.

### Live-cell microscopy

One milliliter of cell culture (6 × 10^6^ cells/mL) were incubated in 1.7 mL Eppendorf tube with 5 μg/mL Hoechst (Cell Signaling Technology) for 2 min and then collected by centrifugation. The culture medium was removed, leaving approximately 10 μL overlaying the cell pellet. Cells were gently resuspended, and 2-3 μL of the suspension was imaged using Axioplan 2 fluorescence microscope (Carl Zeiss Microscopy) equipped with an ORCA-ER camera (Hamamatsu) and AxioVision Rel.4.8 software (Carl Zeiss Microscopy). Exposure times were 200 ms for GFP fluorescence and 50 ms for Hoechst fluorescence and differential interference contrast (DIC). Images were processed using ImageJ version 1.51h (NIH).

### RIP-seq

Cells were induced with 10 μg/mL doxycycline for 5 days to express RBP6 and initiate metacyclic VSG expression. Anti-FLAG M2 magnetic beads (Sigma) were prepared by washing 50 μL of bead slurry twice with 1 mL of wash buffer (PBS containing 0.2% NP-40). Anti-Ty1 beads were prepared by washing 50 μL of Dynabeads Protein G slurry (30 mg/mL; Invitrogen) twice with 1 mL of wash buffer, followed by incubation with approximately 5 μg of BB2 anti-Ty1 monoclonal antibody in 0.5 mL of wash buffer for 2 h at 4°C with gentle rotation. The antibody-coupled beads were subsequently washed five times with 1 mL of wash buffer. Cells expressing Sub2-FLAG (0.5 L culture), TDP1-Ty1, or RBP16-FLAG (1.5 L cultures; 6-8 × 10^6^ cells/mL) were harvested and washed three times with ice-cold PBSG (phosphate-buffered saline containing 25 mM glucose). Cell pellets were lysed in 0.5 mL of ice-cold lysis buffer containing PBS, 1% NP-40, 5% glycerol, 2 mM DTT, 25 mM glucose, 20 mM ribonucleoside vanadyl complex (New England Biolabs), and Complete Mini EDTA-free protease inhibitor cocktail (Roche; one tablet per 5 mL). Following centrifugation at 13,200 rpm for 15 min at 4°C in an Eppendorf 5424 R microcentrifuge, the clarified lysates were transferred to tubes containing the prepared anti-FLAG or anti-Ty1 beads and incubated for 2-3 h at 4°C with gentle rotation. The beads were washed 10 times with 1 mL of ice-cold wash buffer and immunoprecipitated RNA was isolated using TRIzol reagent (Invitrogen). One-fiftieth of each cell lysate was processed directly with TRIzol reagent to generate the corresponding input RNA sample. RNA libraries were prepared from input and immunoprecipitated RNA without poly(A) selection and subjected to Illumina sequenced at the Yale Center for Genome Analysis.

### Sinefungin and A7973 treatments and UV irradiation

Cells were induced with 10 μg/mL doxycycline for 5 days to express RBP6 and initiate differentiation to metacyclic forms. Sinefungin (Enzo Life Sciences) was added to 500-mL cell cultures (8 × 10^6^ cells/mL) to a final concentration of 2 μg/mL. AN7973 (MedChemExpress) was added to parallel cultures to a final concentration of 250 nM. Cultures were incubated at 28°C for 30 min, after which cells were collected by centrifugation and washed three times with ice-cold PBSG (phosphate-buffered saline containing 25 mM glucose) supplemented with the corresponding inhibitor. Cells were resuspended in 1 mL of inhibitor-containing PBSG, transferred to wells of a six-well polystyrene tissue culture plate, placed on ice, and irradiated three times with 300 mJ/cm^2^ UV using an XL-1500 UV crosslinker (Spectronics Corporation). RNA was isolated using TRIzol reagent (Invitrogen). Total RNA and poly(A)+ RNA libraries were prepared and sequenced at the Yale Center for Genome Analysis.

### Bioinformatics

Novaseq pair end 150 bp reads obtained at the Yale Center for Genome Analysis were processed with Galaxy (https://galaxyproject.org/citing-galaxy/)^4^ and reads in FASTA format were mapped to the *T. brucei* 11 megabase chromosomes (GeneDB version 5) or custom-made FASTA files using SeqMan Ngen of the Lasergene v17 software package from DNASTAR. The SeqMan NGen layout algorithm is based on a local match percentage, and the match percentage threshold has to be met in each overlapping window of 50 bases. For the RIP-Seq data we used a minimum aligned length of 75 nt and allowed a maximum of two mismatches. Reads with multiple matches in the genome were distributed randomly between the identical sequences, and the genes were counted as separate. For the RNA-seq data from the effect of UV irradiation, sinefungin/UV, and AN7973/UV on the *T. brucei* transcriptome we used a minimum aligned length of 15 nt. SeqMan Ultra of the Lasergene v17 software package was used to extract read counts from alignments and the Integrative Genomics Viewer^5^ was used to visualize Bam alignments.

For the analysis of SL RNA and mRNA 3′ and 5′ end-containing sequences, a sequence-search code was implemented in Python 3 using only standard-library modules. Target sequences were provided in a CSV file containing numeric sequence identifiers and nucleotide sequences.Input FASTA and FASTQ files were scanned record by record, and each target sequence was searched as an exact substring match against each record sequence. For each input file, the workflow generated one CSV file per target sequence containing matching records, a per-file summary, and a master summary across all processed files. Individual CSV output files were analyzed using TextEdit version 1.20 (Apple).

## REFERENCES

1. Sather, S. and Agabian, N. (1985). A 5’ spliced leader is added in trans to both alpha- and beta-tubulin transcripts in Trypanosoma brucei. Proc. Natl. Acad. Sci. USA. 82, 5695–5699.

2. Sutton, R.E. and Boothroyd, J.C. (1986). Evidence for trans splicing in trypanosomes. Cell 47, 527–535.

3. Matthews, K.R., Tschudi, C., and Ullu, E. (1994). A common pyrimidine-rich motif governs trans-splicing and polyadenylation of tubulin polycistronic pre-mRNA in trypanosomes. Genes Dev. 8, 491–501.

4. Vassella, E., Braun, R., and Roditi, I. (1994). Control of polyadenylation and alternative splicing of transcripts from adjacent genes in a procyclin expression site: a dual role for polypyrimidine tracts in trypanosomes? Nucleic Acids Res. 22, 1359–1364.

5. LeBowitz, J.H., Smith, H.Q., Rusche, L., and Beverley, S.M. (1993). Coupling of poly(A) site selection and trans-splicing in Leishmania. Genes Dev. 7, 996–1007.

6. Chikne, V., Gupta, S.K., Doniger, T., K, S.R., Cohen-Chalamish, S., Waldman Ben-Asher, H., Kolet, L., Yahia, N.H., Unger, R., Ullu, E., Kolev, N.G., Tschudi, C., and Michaeli, S. (2017). The Canonical Poly (A) Polymerase PAP1 Polyadenylates Non-Coding RNAs and Is Essential for snoRNA Biogenesis in Trypanosoma brucei. J. Mol. Biol. 429, 3301–3318.

7. Begolo, D., Vincent, I.M., Giordani, F., Pöhner, I., Witty, M.J., Rowan, T.G., Bengaly, Z., Gillingwater, K., Freund, Y., Wade, R.C., Barrett, M.P., and Clayton, C. (2018). The trypanocidal benzoxaborole AN7973 inhibits trypanosome mRNA processing. PLoS Pathog. 14, e1007315.

8. Koch, H., Raabe, M., Urlaub, H., Bindereif, A., and Preußer, C. (2016). The polyadenylation complex of Trypanosoma brucei: Characterization of the functional poly(A) polymerase. RNA Biol. 13, 221–231.

9. Murphy, W.J., Watkins, K.P., and Agabian, N. (1986). Identification of a novel Y branch structure as an intermediate in trypanosome mRNA processing: evidence for trans splicing. Cell 47, 517–525.

10. Sutton, R.E. and Boothroyd, J.C. (1988). Trypanosome trans-splicing utilizes 2’-5’ branches and a corresponding debranching activity. EMBO J. 7, 1431–1437.

11. Tschudi, C., Richards, F.F., and Ullu, E. (1986). The U2 RNA analogue of Trypanosoma brucei gambiense: implications for a splicing mechanism in trypanosomes. Nucleic Acids Res. 14, 8893–8903.

12. Mottram, J., Perry, K.L., Lizardi, P.M., Lührmann, R., Agabian, N., and Nelson, R.G. (1989). Isolation and sequence of four small nuclear U RNA genes of Trypanosoma brucei subsp. brucei: identification of the U2, U4, and U6 RNA analogs. Mol. Cell. Biol. 9, 1212–1223.

13. Tschudi, C., Williams, S.P., and Ullu, E. (1990). Conserved sequences in the U2 snRNA-encoding genes of Kinetoplastida do not include the putative branchpoint recognition region. Gene 91, 71–77.

14. Miller, S.I., and Wirth, D.F. (1988). trans splicing in Leishmania enriettii and identification of ribonucleoprotein complexes containing the spliced leader and U2 equivalent RNAs. Mol. Cell. Biol. 8, 2597–2603.

15. Hartshorne, T., and Agabian, N. (1990). A new U2 RNA secondary structure provided by phylogenetic analysis of trypanosomatid U2 RNAs. Genes Dev. 4, 2121–2131.

16. Patzelt, E., Perry, K.L., and Agabian, N. (1989). Mapping of branch sites in trans-spliced pre-mRNAs of Trypanosoma brucei. Mol. Cell. Biol. 9, 4291–4297.

17. Lücke, S., Jürchott, K., Hung, L.H., and Bindereif, A. (2005). mRNA splicing in Trypanosoma brucei: branch-point mapping reveals differences from the canonical U2 snRNA-mediated recognition. Mol. Biochem. Parasitol. 142, 248–251.

18. Preußer, C., Rossbach, O., Hung, L.H., Li, D., and Bindereif A. (2014). Genome-wide RNA-binding analysis of the trypanosome U1 snRNP proteins U1C and U1-70K reveals cis/trans-spliceosomal network. Nucleic Acids Res. 42, 6603–6615.

19. Tkacz, I.D., Lustig, Y., Stern, M.Z., Biton, M., Salmon-Divon, M., Das, A., Bellofatto, V., and Michaeli, S. (2007). Identification of novel snRNA-specific Sm proteins that bind selectively to U2 and U4 snRNAs in Trypanosoma brucei. RNA 13, 30–43.

20. Palfi, Z., Jaé, N., Preusser, C., Kaminska, K.H., Bujnicki, J.M., Lee, J.H., Günzl, A., Kambach, C., Urlaub, H., and Bindereif, A. (2009). SMN-assisted assembly of snRNP-specific Sm cores in trypanosomes. Genes Dev. 23, 1650–1664.

21. Yong, J., Wan, L., and Dreyfuss, G. (2004). Why do cells need an assembly machine for RNA-protein complexes? Trends Cell Biol. 14, 226–232.

22. Bangs, J.D., Crain, P.F., Hashizume, T., McCloskey, J.A., and Boothroyd, J.C. (1992). Mass spectrometry of mRNA cap 4 from trypanosomatids reveals two novel nucleosides. J. Biol. Chem. 267, 9805–9815.

23. Liang, X.H., Xu, Y.X., and Michaeli, S. (2002). The spliced leader-associated RNA is a trypanosome-specific sn(o) RNA that has the potential to guide pseudouridine formation on the SL RNA. RNA 8, 237–246.

24. Perry, K.L., Watkins, K.P., and Agabian, N. (1987). Trypanosome mRNAs have unusual “cap 4” structures acquired by addition of a spliced leader. Proc. Natl. Acad. Sci. USA 84, 8190–8194.

25. Freistadt, M.S., Cross, G.A., Branch, A.D., and Robertson, H.D. (1987). Direct analysis of the mini-exon donor RNA of Trypanosoma brucei: detection of a novel cap structure also present in messenger RNA. Nucleic Acids Res. 15, 9861–9879.

26. Bangs, J.D. (2018). Evolution of Antigenic Variation in African Trypanosomes: Variant Surface Glycoprotein Expression, Structure, and Function. Bioessays 40, e1800181.

27. Horn, D. (2014). Antigenic variation in African trypanosomes. Mol. Biochem. Parasitol. 195, 123–129.

28. Navarro, M., and Gull, K. (2001). A pol I transcriptional body associated with VSG mono-allelic expression in Trypanosoma brucei. Nature 414, 759–763.

29. López-Escobar, L., Hänisch, B., Halliday, C., Ishii, M., Akiyoshi, B., Dean, S., Sunter, J.D., Wheeler, R.J., and Gull, K. (2022). Stage-specific transcription activator ESB1 regulates monoallelic antigen expression in Trypanosoma brucei. Nat. Microbiol. 7, 1280–1290.

30. Faria, J., Luzak, V., Müller, L.S.M., Brink, B.G., Hutchinson, S., Glover, L., Horn, D., and Siegel, T.N. (2021). Spatial integration of transcription and splicing in a dedicated compartment sustains monogenic antigen expression in African trypanosomes. Nat. Microbiol. 6, 289–300.

31. Budzak, J., Jones, R., Tschudi, C., Kolev, N.G., and Rudenko, G. (2022). An assembly of nuclear bodies associates with the active VSG expression site in African trypanosomes. Nat. Commun. 13, 101.

32. Ridewood, S., Ooi, C.P., Hall, B., Trenaman, A., Wand, N.V., Sioutas, G., Scherwitzl, I., and Rudenko, G. (2017). The role of genomic location and flanking 3’UTR in the generation of functional levels of variant surface glycoprotein in Trypanosoma brucei. Mol. Microbiol. 106, 614–634.

33. Bakari-Soale, M., Batram, C., Zimmermann, H., Jones, N.G., and Engstler, M. (2026). The Dual Role of the 16mer Motif Within the 3’ Untranslated Region of the Variant Surface Glycoprotein of Trypanosoma brucei. Mol. Microbiol. 125, 67–79.

34. Melo do Nascimento, L., Egler, F., Arnold, K., Papavasiliou, N., Clayton, C., and Erben E. (2021). Functional insights from a surface antigen mRNA-bound proteome. Elife 10, e68136.

35. Bravo Ruiz, G., Tinti, M., Ridgway, M., and Horn, D. (2022). Control of Variant Surface Glycoprotein Expression by CFB2 in Trypanosoma brucei and Quantitative Proteomic Connections to Translation and Cytokinesis. mSphere 7, e0006922.

36. Amiguet-Vercher, A., Pérez-Morga, D., Pays, A., Poelvoorde, P., Van Xong, H., Tebabi, P., Vanhamme, L., and Pays, E. (2004). Loss of the mono-allelic control of the VSG expression sites during the development of Trypanosoma brucei in the bloodstream. Mol. Microbiol. 51, 1577–1588.

37. Pays, E., Tebabi, P., Pays, A., Coquelet, H., Revelard, P., Salmon, D., and Steinert, M. (1989). The genes and transcripts of an antigen gene expression site from T. brucei. Cell 57, 835–845.

38. Kolev, N.G., Ramsdell, T.K., and Tschudi, C. (2018). Temperature shift activates bloodstream VSG expression site promoters in Trypanosoma brucei. Mol. Biochem. Parasitol. 226, 20–23.

39. Kolev, N.G., Ramey-Butler, K., Cross, G.A., Ullu, E., and Tschudi, C. (2012). Developmental progression to infectivity in Trypanosoma brucei triggered by an RNA-binding protein. Science 338, 1352–1353.

40. Toh, J.Y., Nkouawa, A., Dong, G., Kolev, N.G., Tschudi, C. (2023). Two cold shock domain containing proteins trigger the development of infectious Trypanosoma brucei. PLoS Pathog. 19, e1011438.

41. Narayanan, M.S., and Rudenko, G. (2013). TDP1 is an HMG chromatin protein facilitating RNA polymerase I transcription in African trypanosomes. Nucleic Acids Res. 41, 2981–2992.

42. Kosugi, S., Hasebe, M., Tomita, M., and Yanagawa, H. (2008). Nuclear export signal consensus sequences defined using a localization-based yeast selection system. Traffic 9, 2053–2062.

43. Bühlmann, M., Walrad, P., Rico, E., Ivens, A., Capewell, P., Naguleswaran, A., Roditi, I., and Matthews, K.R. (2015). NMD3 regulates both mRNA and rRNA nuclear export in African trypanosomes via an XPOI-linked pathway. Nucleic Acids Res. 43, 4491–4504.

44. Marchetti, M.A., Tschudi, C., Kwon, H., Wolin, S.L., and Ullu, E. (2000). Import of proteins into the trypanosome nucleus and their distribution at karyokinesis. J. Cell Sci. 113, 899–906.

45. Bradatsch, B., Katahira, J., Kowalinski, E., Bange, G., Yao, W., Sekimoto, T., Baumgärtel, V., Boese, G., Bassler, J., Wild, K., Peters, R., Yoneda, Y., Sinning, I., and Hurt, E. (2007). Arx1 functions as an unorthodox nuclear export receptor for the 60S preribosomal subunit. Mol. Cell 27, 767–779.

46. Hung, N.J., Lo, K.Y., Patel, S.S., Helmke, K., and Johnson, A.W. (2008). Arx1 is a nuclear export receptor for the 60S ribosomal subunit in yeast. Mol. Biol. Cell 19, 735–744.

47. Florini, F., Naguleswaran, A., Gharib, W.H., Bringaud, F., and Roditi, I. (2019). Unexpected diversity in eukaryotic transcription revealed by the retrotransposon hotspot family of Trypanosoma brucei. Nucleic Acids Res. 47, 1725–1739.

48. Pandey, P., Wackowski, K., Dubey, A.P., and Read, L.K. (2025). DRBD18 acts as a transcript-specific RNA editing auxiliary factor in Trypanosoma brucei. RNA 31, 245–257.

49. Bishola Tshitenge, T., and Clayton, C. (2022). The Trypanosoma brucei RNA-binding protein DRBD18 ensures correct mRNA trans splicing and polyadenylation patterns. RNA 28, 1239–1262.

50. Bard, J.E., Tylec, B.L., Dubey, A.P., Lamb, N.A., Yergeau, D.A., and Read, L.K. (2024). Life stage-specific poly(A) site selection regulated by Trypanosoma brucei DRBD18. Proc. Natl. Acad. Sci. USA 121, e2403188121.

51. Mishra, A., Kaur, J.N., McSkimming, D.I., Hegedűsová, E., Dubey, A.P., Ciganda, M., Paris, Z., and Read, L.K. (2021). Selective nuclear export of mRNAs is promoted by DRBD18 in Trypanosoma brucei. Mol. Microbiol. 116, 827–840.

52. Kramer, S. (2021). Nuclear mRNA maturation and mRNA export control: from trypanosomes to opisthokonts. Parasitology 148, 1196–1218.

53. Pelletier, M., and Read, L.K. (2003). RBP16 is a multifunctional gene regulatory protein involved in editing and stabilization of specific mitochondrial mRNAs in Trypanosoma brucei. RNA 9, 457–468.

54. Goos, C., Dejung, M., Janzen, C.J., Butter, F., and Kramer, S. (2017). The nuclear proteome of Trypanosoma brucei. PLoS One 12, e0181884.

55. Patrick, K.L., Shi, H., Kolev, N.G., Ersfeld, K., Tschudi, C., and Ullu, E. (2009). Distinct and overlapping roles for two Dicer-like proteins in the RNA interference pathways of the ancient eukaryote Trypanosoma brucei. Proc. Natl. Acad. Sci. USA 106, 17933–17938.

56. Hung, M.L., Hautbergue, G.M., Snijders, A.P., Dickman, M.J., and Wilson, S.A. (2010). Arginine methylation of REF/ALY promotes efficient handover of mRNA to TAP/NXF1. Nucleic Acids Res. 38, 3351–3361.

57. Goulah, C.C., Pelletier, M., Read, L.K. (2006). Arginine methylation regulates mitochondrial gene expression in Trypanosoma brucei through multiple effector proteins. RNA 12, 1545–1555.

58. Lott, K., Mukhopadhyay, S., Li, J., Wang, J., Yao, J., Sun, Y., Qu, J., and Read, L.K. (2015). Arginine methylation of DRBD18 differentially impacts its opposing effects on the trypanosome transcriptome. Nucleic Acids Res. 43, 5501–5523.

59. Brahms, H., Meheus, L., de Brabandere, V., Fischer, U., and Lührmann, R. (2001). Symmetrical dimethylation of arginine residues in spliceosomal Sm protein B/B’ and the Sm-like protein LSm4, and their interaction with the SMN protein. RNA 7, 1531–1542.

60. Fleckner, J., Zhang, M., Valcárcel, J., Green, M.R. (1997). U2AF65 recruits a novel human DEAD box protein required for the U2 snRNP-branchpoint interaction. Genes Dev. 11, 1864–1872.

61. Shen, H., Zheng, X., Shen, J., Zhang, L., Zhao, R., and Green, M.R. (2008). Distinct activities of the DExD/H-box splicing factor hUAP56 facilitate stepwise assembly of the spliceosome. Genes Dev. 22, 1796–1803.

62. Luo, M.L., Zhou, Z., Magni, K., Christoforides, C., Rappsilber, J., Mann, M., and Reed, R. (2001). Pre-mRNA splicing and mRNA export linked by direct interactions between UAP56 and Aly. Nature 413, 644–647.

63. Strässer, K., and Hurt, E. (2001). Splicing factor Sub2p is required for nuclear mRNA export through its interaction with Yra1p. Nature 413, 648–652.

64. Hohmann, U., Graf, M., Tirián, L., Pacheco-Fiallos, B., Schellhaas, U., Fin, L., Handler, D., Phillips, A.W., Riabov-Bassat, D., Faraway, R., Pühringer, T., Szalay, M.F., Roitinger, E., Brennecke, J., and Plaschka, C. (2026). An ATP-gated molecular switch orchestrates human mRNA export. Nature 649, 1042–1050.

65. Serpeloni, M., Moraes, C.B., Muniz, J.R., Motta, M.C., Ramos, A.S., Kessler, R.L., Inoue, A.H., daRocha, W.D., Yamada-Ogatta, S.F., Fragoso, S.P., Goldenberg, S., Freitas-Junior, L.H., and Avila, A.R. (2011). An essential nuclear protein in trypanosomes is a component of mRNA transcription/export pathway. PLoS One 6, 20730.

66. Watkins, K.P., and Agabian, N. (1991). In vivo UV cross-linking of U snRNAs that participate in trypanosome trans-splicing. Genes Dev. 5,1859–1869.

67. Mandel, C.R., Kaneko, S., Zhang, H., Gebauer, D., Vethantham, V., Manley, J.L., and Tong, L. (2006). Polyadenylation factor CPSF-73 is the pre-mRNA 3’-end-processing endonuclease. Nature 444, 953–956.

68. Schönemann, L., Kühn, U., Martin, G., Schäfer, P., Gruber, A.R., Keller, W., Zavolan, M., and Wahle, E. (2014). Reconstitution of CPSF active in polyadenylation: recognition of the polyadenylation signal by WDR33. Genes Dev. 28, 2381–2393.

69. Shi, Y., and Manley, J.L. (2015). The end of the message: multiple protein-RNA interactions define the mRNA polyadenylation site. Genes Dev. 29, 889–897.

70. Lerner, M.R., Boyle, J.A., Mount, S.M., Wolin, S.L., and Steitz, J.A. (1980). Are snRNPs involved in splicing? Nature 283, 220–224.

71. Ammerman, M.L., Fisk, J.C., and Read, L.K. (2008). gRNA/pre-mRNA annealing and RNA chaperone activities of RBP16. RNA 14, 1069–1080.

72. Dai, L., Chai, D., Gu, S.Q., Gabel, J., Noskov, S.Y., Blocker, F.J., Lambowitz, A.M., and Zimmerly, S. (2008). A three-dimensional model of a group II intron RNA and its interaction with the intron-encoded reverse transcriptase. Mol. Cell 30, 472–485.

73. Hilliker, A.K., and Staley, J.P. (2004). Multiple functions for the invariant AGC triad of U6 snRNA. RNA 10, 921–928.

74. Datta, B., and Weiner, A.M. (1993). The phylogenetically invariant ACAGAGA and AGC sequences of U6 small nuclear RNA are more tolerant of mutation in human cells than in Saccharomyces cerevisiae. Mol. Cell Biol. 13, 5377–5382.

75. Henze, K., Badr, A., Wettern, M., Cerff, R., and Martin, W. (1995). A nuclear gene of eubacterial origin in Euglena gracilis reflects cryptic endosymbioses during protist evolution. Proc. Natl. Acad. Sci. USA. 92, 9122–9126.

76. Thakur, P.K., Butenko, A., Karásek, F., Svobodová, M., Faktorová, D., Pavlisková, H., Varga, V., Horák, A., Lukeš, J., and Staněk, D. (2025). Splice site diversity and abundance of noncanonical introns in diplonemids (Diplonemea, Euglenozoa). RNA 31, 1826–1840.

77. Kolev, N.G., Rajan, K.S., Tycowski, K.T., Toh, J.Y., Shi, H., Lei, Y., Michaeli, S., and Tschudi, C. (2019). The vault RNA of Trypanosoma brucei plays a role in the production of trans-spliced mRNA. J Biol. Chem. 294, 15559–15574.

78. Ares, M.Jr., Igel, H., Katzman, S., and Donohue, J.P. (2024). Intron lariat spliceosomes convert lariats to true circles: implications for intron transposition. Genes Dev. 38, 322–335.

79. McNally, K.P., and Agabian, N. (1992). Trypanosoma brucei spliced-leader RNA methylations are required for trans splicing in vivo. Mol. Cell. Biol. 12, 4844–4851.

80. Watkins, K.P., Dungan, J.M., and Agabian, N. (1994). Identification of a small RNA that interacts with the 5’ splice site of the Trypanosoma brucei spliced leader RNA in vivo. Cell 76, 171–182.

81. Cohen, S.B., Graham, M.E., Lovrecz, G.O., Bache, N., Robinson, P.J., and Reddel, R.R. (2007). Protein composition of catalytically active human telomerase from immortal cells. Science 315, 1850–1853.

82. Tarn, W.Y., and Steitz, J.A. (1996). A novel spliceosome containing U11, U12, and U5 snRNPs excises a minor class (AT-AC) intron in vitro. Cell 84, 801–811.

83. Kaida, D., Berg, M.G., Younis, I., Kasim, M., Singh, L.N., Wan, L., and Dreyfuss, G. (2010). U1 snRNP protects pre-mRNAs from premature cleavage and polyadenylation. Nature 468, 664–668.

84. Berg, M.G., Singh, L.N., Younis, I., Liu, Q., Pinto, A.M., Kaida, D., Zhang, Z., Cho, S., Sherrill-Mix, S., Wan, L., and Dreyfuss, G. (2012). U1 snRNP determines mRNA length and regulates isoform expression. Cell 150, 53–64.

85. So, B.R., Di, C., Cai, Z., Venters, C.C., Guo, J., Oh, J.M., Arai, C., and Dreyfuss, G. (2019). A Complex of U1 snRNP with Cleavage and Polyadenylation Factors Controls Telescripting, Regulating mRNA Transcription in Human Cells. Mol. Cell 76, 590–599.

86. Mowry, K.L., and Steitz, J.A. (1987). Identification of the human U7 snRNP as one of several factors involved in the 3’ end maturation of histone premessenger RNA’s. Science 238, 1682–1687.

87. Box, J.A., Bunch, J.T., Tang, W., and Baumann, P. (2008). Spliceosomal cleavage generates the 3’ end of telomerase RNA. Nature 456, 910–914.

88. Kannan, R., Helston, R.M., Dannebaum, R.O., and Baumann, P. (2015). Diverse mechanisms for spliceosome-mediated 3’ end processing of telomerase RNA. Nat. Commun. 6, 6104.

89. Qi, X., Rand, D.P., Podlevsky, J.D., Li, Y., Mosig, A., Stadler, P.F., and Chen, J.J. (2015). Prevalent and distinct spliceosomal 3’-end processing mechanisms for fungal telomerase RNA. Nat. Commun. 6, 6105.

90. Baillat, D., Hakimi, M.A., Näär, A.M., Shilatifard, A., Cooch, N., and Shiekhattar, R. (2005). Integrator, a multiprotein mediator of small nuclear RNA processing, associates with the C-terminal repeat of RNA polymerase II. Cell 123, 265–276.

91. Pfleiderer, M.M., and Galej, W.P. (2021). Structure of the catalytic core of the Integrator complex. Mol. Cell 81, 1246–1259.

## Methods References

1. Frejno, M., Berger, M.T., Tüshaus, J. et al. (2025). Unifying the analysis of bottom-up proteomics data with CHIMERYS. Nat. Methods 22, 1017–1027.

2. Zolg, D.P., Gessulat, S., Paschke, C. et al. (2021). INFERYS rescoring: Boosting peptide identifications and scoring confidence of database search results. Rapid Commun. Mass Spectrom. 39, e9128.

3. Käll, L., Canterbury, J.D., Weston, J. et al. (2007). Semi-supervised learning for peptide identification from shotgun proteomics datasets. Nat. Methods 4, 923–925.

4. Galaxy Community. (2022). The Galaxy platform for accessible, reproducible and collaborative biomedical analyses: 2022 update. Nucleic Acids Res. 50, W345–W351.

5. Robinson, J.T., Thorvaldsdóttir, H., Winckler, W., Guttman, M., Lander, E.S., Getz, G., and Mesirov, J.P. (2011). Integrative Genomics Viewer. Nat. Biotechnology 29, 24–26.

