## Supplementary material for "A *trans-*splicing model for poly(A)-tail addition during VSG mRNA maturation in *Trypanosoma brucei*": Figures S1-S10

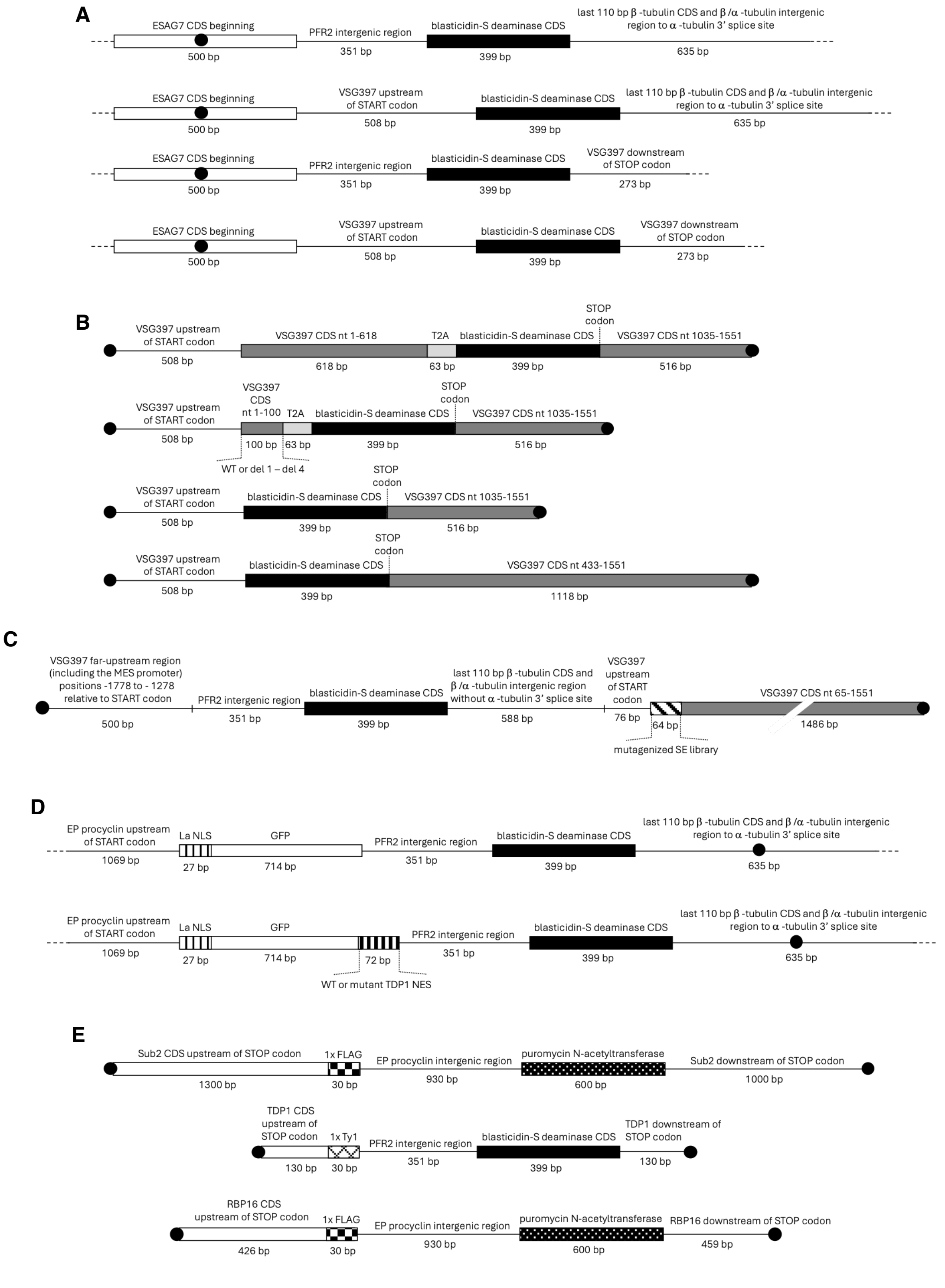


**Figure S1. Schematic representation of the constructs used in this study**

The organization of reporter constructs, mutagenized libraries, and expression cassettes used for the analyses described in this study is shown (not to scale). Black circles indicate linearization points prior to transfection.

(A) Constructs used for testing the VSG397 pre-mRNA SL-addition site strength.

(B) Deletion constructs for the identification of the VSG397 CDS splicing enhancer. Note the use of T2A peptide [Tran, K.D., Rodriguez-Contreras, D., Vieira, D.P., Yates, P.A., David, L., Beatty, W., Elferich, J., and Landfear, S.M. (2013). KHARON1 mediates flagellar targeting of a glucose transporter in *Leishmania mexicana* and is critical for viability of infectious intracellular amastigotes. J. Biol. Chem. *288*, 22721-22733] for introducing a polypeptide chain discontinuity between the N-terminus of VSG397 (bearing a signal peptide) and the blasticidin-S deaminase CDS.

(C) Doped mutagenesis library construct for identifying nucoetides important for the function of the splicing enhancer.

(D) GFP-fusion assay constructs for testing the functionality of the putative TDP1 nuclear export signal.

(E) Epitope-tagging constructs for Sub2, TDP1, and RBP16.

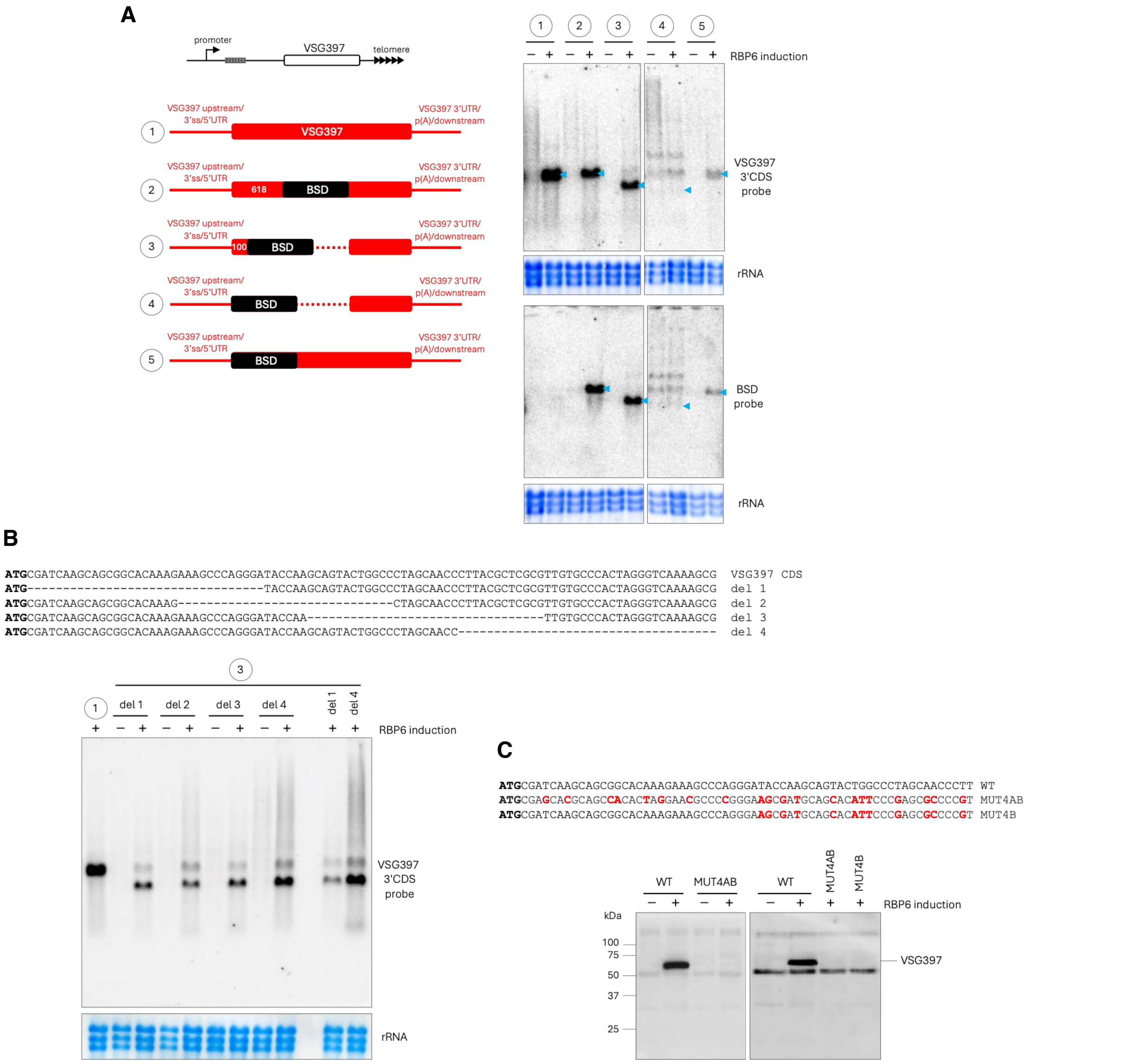


**Figure S2. Deletion analysis defines the splicing enhancer at the beginning of the VSG397 CDS**

(A) Reporter constructs used to assess the effect of VSG397 CDS deletions on mRNA production. A schematic representation of the endogenous metacyclic-form VSG397 expression site (the construct integration site) is shown above. Northern blot analyses of reporter mRNA abundance in procyclic cells induced to differentiate into metacyclic forms for 5 days through RBP6 expression are shown on the right. Blue arrowheads indicate the expected sizes of properly matured mRNAs for each construct.

(B) Deletion analysis of the splicing enhancer within the VSG397 CDS. The sequences of the tested constructs and northern blot analyses of their expression levels after 5 days of RBP6-induced expression are shown.

(C) Assessment of the suitability of a sequence identified in the doped mutagenesis library screen as a control for the protein pulldown assay. The sequences of the tested constructs are shown. Western blotting with a rabbit polyclonal anti-VSG397 antibody was used to determine expression levels.

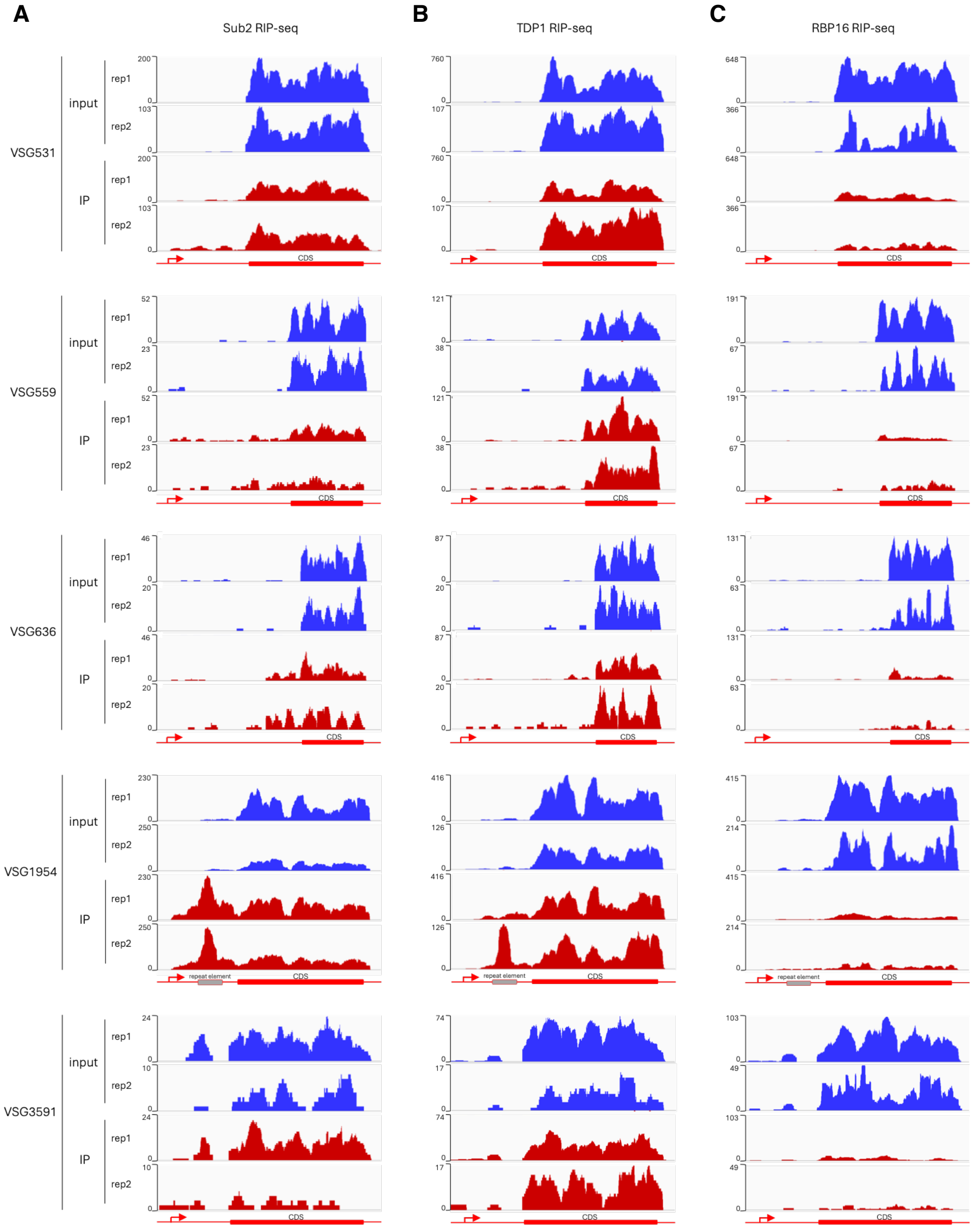


**Figure S3. RIP-seq analysis of Sub2 (A), TDP1 (B), and RBP16 (C) at the remaining five metacyclic VSG expression sites in *T. brucei* Lister 427.**
Data from two biological replicates (rep1 and rep2) are shown. IP, immunoprecipitate.

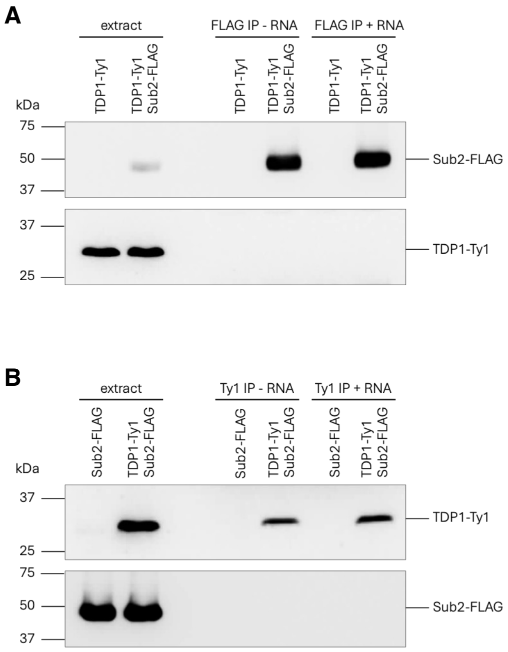


**Figure S4. Sub2 and TDP1 do not co-immunoprecipitate**
(A) Anti-FLAG immunoprecipitation of Sub2-FLAG from extracts prepared from cells expressing the indicated endogenously tagged constructs, performed in the presence or absence of the wild-type 15-nt bait RNA. Western blotting was used to detect FLAG (top) and Ty1 (bottom).
(B) Reciprocal anti-Ty1 (BB2 antibody) immunoprecipitation of TDP1-Ty1 from extracts prepared from cells expressing the indicated endogenously tagged constructs, performed in the presence or absence of the wild-type 15-nt bait RNA. Western blotting was used to detect Ty1 (top) and FLAG (bottom).

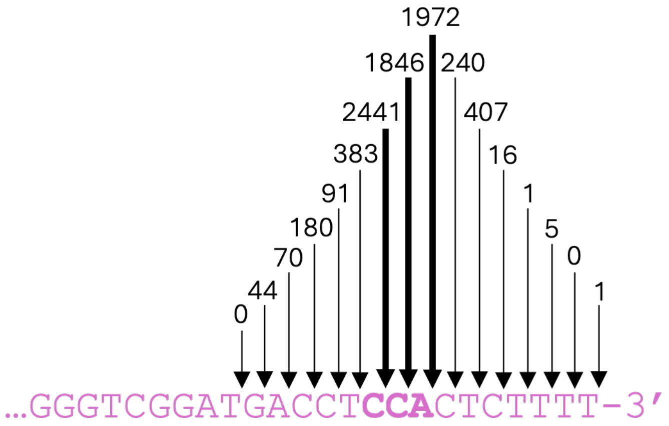


**Figure S5. Distribution of U1C iCLIP reads terminating at the 3′ end of SL RNA**
Number of U1C iCLIP reads (raw data from Preußer *et al.*, 2014)[18] terminating at the indicated positions at the 3′ end of SL RNA.

```
                                                                      catalytic
                                                        SL 5′ ss      triad
                          branch site
                          adenosine         SL exon                          SL intron
Blastocrithidia gerricola     AACTAACGCTATATAA-GTATCAGTTTCTGTACTTTATTGGTATGAGAAGCTTCCGGATCCATCAA-
Blastocrithidia leptocoridis  AACTAACGCATTTT-TTGTTACAGTTTCTGTACTTTATTGGTATGAGAAACTTCCAGAACATTCAT-
Blastocrithidia miridarum     AACTAACGCTATATAA-GTATCAGTTTCTGTACTTTATTGGTATGAGAAGCTTCCGGATCCATCAA-
Blastocrithidia triatomae     AACTAACGCATTTT-TGGTTACAGTTTCTGTACTTTATTGGTATGAGAAACTTCCAGAACTTTAGT-
Bodo saltans                  AACTAACGCTATAAAA-GATACAGTTTCTGTACTATATTGGTATGAGGAACTTTCCAGTTCTAAAC-
Crithidia luciliae            AACTAACGCTATATAA-GTATCAGTTTCTGTACTTTATTGGTATAAGAAACTTCCGGTCCACGCCG-
Crithidia confusa             AACTAACGCTATATAA-GTATCAGTTTCTGTACTATATTGGTATGTGAAGCCATCCGTGCCAAGTG-
Crithidia insperata           AACTAACGCTATATAA-GTATCAGTTTCTGTACTATATTGGTATGAGAAGCTTCCGGAATCCTCTT-
Crithidia abscondita          AACTAACGCTATATAA-GTATCAGTTTCTGTACTATATTGGTATATGAAGCTTCCGGTAACTGTAA-
Crithidia brevicula           AACTAACGCTATATAA-GTATCAGTTTCTGTACTTTATTGGTATGAGAAGCTTCCGGATCCATCAA-
Crithidia deanei              AACTAAAGCATTATATAGATACAGTTTCTGTACTATATTGGTATGTGAAACTTCCAAGTCTAACTT-
Cryptobia helicis             AACTTACGCTATAAAA-GTCACAGTTTCTGTACTTTATTGGTATGAGAAGCTTTCCAGCAATGGAA-
Herpetomonas muscarum         AACTAACGCTAAAAATTGTTACAGTTTCTGTACTATATTGGTATGAGAAGCTTCCAGTTCATCTGG-
Herpetomonas megaseliae       AACTAACGCTAAAAATTGTTACAGTTTCTGTACTCTATTGGTATGAGAAGCTTCCAGTTCATCTGG-
Herpetomonas roitmani         AACTAAAGCATTATATAGATACAGTTTCTGTACTTTATTGGTATGTGAAACTTCCAAGTCTAACTT-
Herpetomonas mariadeanei      AACTAACGCTAT-TATTGTTACAGTTTCTGTACTTTATTGGTATGGGAAGCTTCCAGAACATCTAG-
Leishmania tarentolae         AACTAACGCTATATAA-GTATCAGTTTCTGTACTTTATTGGTATGCGAAACTTCCGGAACCTGTCT-
Leishmania hoogstraali        AACTAACGCTATATAA-GTATCAGTTTCTGTACTTTATTGGTATGCGAAACTTCCGGAACCTGTCT-
Leishmania adleri             AACTAACGCTATATAA-GTATCAGTTTCTGTACTTTATTGGTATGCGAAACTTCCGGAACCCGTCT-
Leishmania gymnodactyli       AACTAACGCTATATAA-GTATCAGTTTCTGTACTTTATTGGTATGCGAAACTTCCGGAACCTGTCT-
Leptomonas scantii            AACTAACGCTATATAA-GTATCAGTTTCTGTACTTTATTGGTATGAGAAGCTTCCGGGACCCGTAT-
Leptomonas pyrrhocoris        AACTAACGCTATATAA-GTATCAGTTTCTGTACTTTATTGGTACGTGAAACTTCCGGTATCCCACC-
Leptomonas tenua              AACTAACGCTATATAA-GTATCAGTTTCTGTACTATATTGGTATGAGAAGCTTCCGGATCACGTTA-
Leptomonas jaderae            AACTAACGCTATATAA-GTATCAGTTTCTGTACTATATTGGTATGAGAAGCTTCCGGAACCAGTAT-
Leptomonas neopamerae         AACTAACGCTATATAA-GTATCAGTTTCTGTACTTTATTGGTATGAGAAGCTTCCGGAATCTCGAT-
Leptomonas bifurcata          AACTAACGCTATATAA-GTATCAGTTTCTGTACTATATTGGTATGAGAAGCTTCCGGAAACATCAT-
Leptomonas peterhoffi         AACTAACGCTATATAA-GTATCAGTTTCTGTACTTTATTGGTATGAGAAGCTTCCGGATCCATCAA-
Leptomonas podlipaevi         AACTAACGCTATATAA-GTATCAGTTTCTGTACTATATTGGTATGAGAAGCTTCCGGAACCTATAT-
Leptomonas tarcoles           AACTAACGCTATATAA-GTATCAGTTTCTGTACTTTATTGGTATAAGAAGCTTCCGGTCCACACCG-
Leptomonas costaricensis      AACTAACGCTATATAA-GTATCAGTTTCTGTACTATATTGGTATGCGAAACTTCCGGAACCCTCTT-
Leptomonas spiculata          AACTAACGCTATATAA-GTATCAGTTTCTGTACTATATTGGTATAAGAAGCTTCCGGTCCACACCG-
Phytomonas serpens            AACTAACGCTAT-TCTAGATACAGTTTCTGTACTTTATTGGTATGAGAAACTTCCAGAATTCACGT-
Trypanosoma conorhini         AACTAACGCTAT-TATTGATACAGTTTCTGTACTATATTGGTATGCAGCGCTTCCGGATGCGTCCG-
Trypanosoma dionisii          AACTAACGCTAT-TATTGATACAGTTTCTGTACTATATTGGTACGCGAAGCTTCCGGGGGCGGGCC-
Trypanosoma leeuwenhoeki      AACTAACGCTAT-TATTGATACAGTTTCTGTACTATATTGGTATGCAGCGCTTCCGGATGCGTCCG-
Trypanosoma lewisi            AACTAACGCTAT-TATTGATACAGTTTCTGTACTATATTGGTATGAAAAGCTTCCGGATCCGCCGC-
Trypanosoma mega              AACTAACGCTAT-TATTGATACAGTTTCTGTACTATATTGGTATGAGAAGCTTCCAGAACATCAAG-
Trypanosoma microti           AACTAACGCTAT-TATTGATACAGTTTCTGTACTATATTGGTATGAGAAGCTTCCGGGTCCGCCGC-
Trypanosoma rangeli           AACTAACGCTAT-TATTGATACAGTTTCTGTACTATATTGGTATGCAGCGCTTCCGGATGCGTCCG-
Trypanosoma theileri          AACTAACGATAT-TATTGATACAGTTTCTGTACTATATTGGTATGAGAAGCTTCCAGGAACTTCCT-
Trypanosoma vivax             AACTAAAGCTTTTATTAGA-ACAGTTTCTGTACTATATTGGTATGAGAAGCTCCCGGTCGCAAGAC-
Trypanosoma grayi             AACTAACGCTAT-TATTGATACAGTTTCTGTACTATATTGGTATGAGAAGCTTCCGGAATTGCCAA-
Trypanosoma danilewskyi       AACTAAAGTTAT-TATTGATACAGTTTCTGTACTATATTGGTATGAGAAACTTCCAGGCTAATTAA-
Trypanosoma cobitis           AACTAAAGTTAT-TATTGATACAGTTTCTGTACTATATTGGTATGAGAAACTTCCAGGCTTATTAT-
Trypanosoma cyclops           AACTAAAGATTT-TATTGTTACAGTTTCTGTACTATATTGGTATGCGAAGCTTCCGGGGACATCCG-
Trypanosoma simiae            AACTAAAATTATTTATA-ATACAGTTTCTGTACTATATTGGTATGAGAAGCTCCCAGGACACAAAT-
Trypanosoma avium             AACTAAAGCTATTATTAGA-ACAGTTTCTGTACTATATTGGTATGAGAAGCTTCCAGAATCATCAA-
Trypanosoma cruzi             AACTAACGCTAT-TATTGATACAGTTTCTGTACTATATTGGTACGCGAAGCTTCCAAATCCGCGAG-
Trypanosoma brucei            AACTAACGCTATTATTAGA-ACAGTTTCTGTACTATATTGGTATGAGAAGCTCCCAGTAGCAGCTG-
```

**Figure S6. Alignment of the 5′ regions of trypanosomatid SL RNA sequences**
The intron sequences shown are not aligned. Most of the sequences were obtained from Krchňáková, Z., Krajčovič, J., and Vesteg, M. (2017). On the possibility of an early evolutionary origin for the spliced leader *trans*-splicing. J. Mol. Evol. *85*, 37-45.

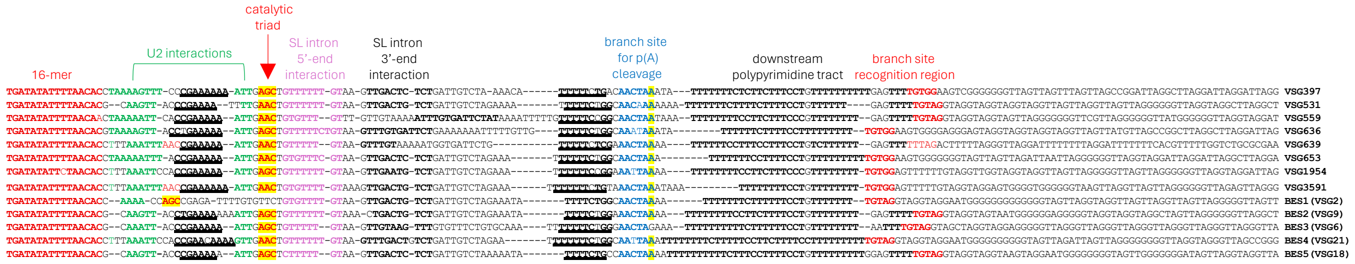


**Figure S7. Sequence alignment of the VSG 16-mer and downstream regions from all eight metacyclic VSGs and VSGs from five bloodstream expression sites (BESs) in *T. brucei* Lister 427**

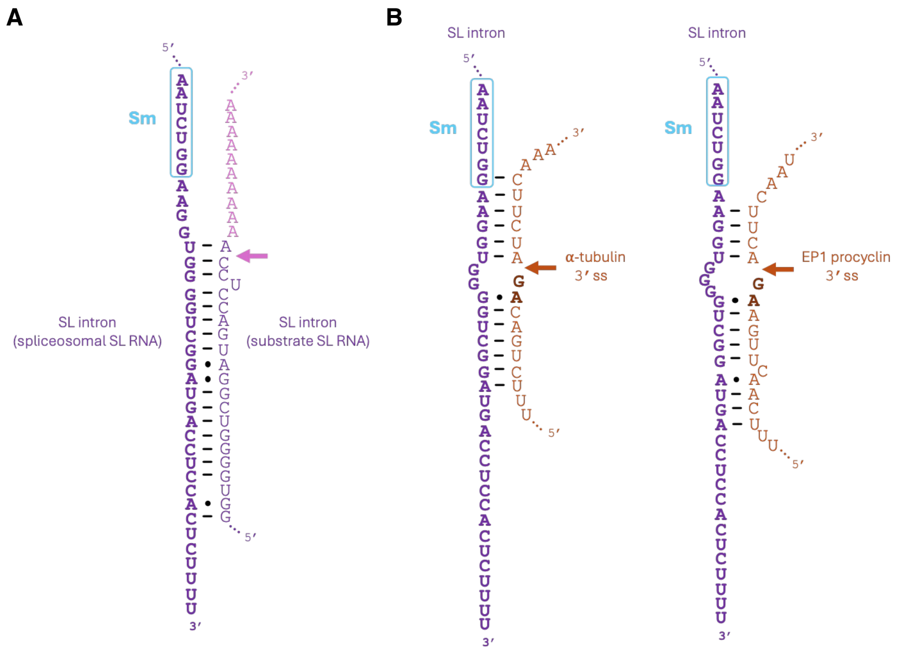


**Figure S8. Potential base-pairing interactions involved in the second step of general (non-VSG) poly(A)-tail and SL *trans*-splicing**

(A) Predicted interaction between the intron of the spliceosomal SL RNA and the polyadenylated substrate SL RNA.

(B) Predicted base-pairing interactions between α-tubulin (left) and EP1 procyclin (right) pre-mRNAs and the intron of the spliceosomal SL RNA.

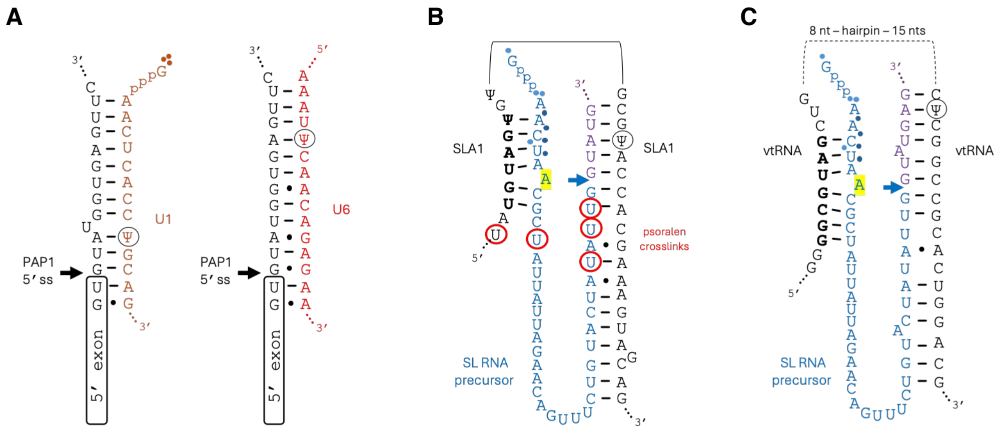


**Figure S9. Potential 5′ splice-site recognition interactions in *cis*- and *trans*-spliceosomes of *T. brucei***

(A) Recognition of the PAP1 intron 5′ splice site by U1 and U6 snRNAs during *cis*-splicing.

(B) Predicted base-pairing interactions between SLA1 RNA and the SL RNA 5′ splice site and branch-site sequence during SL *trans*-splicing. Previously identified psoralen cross-links[80] between a single uridine in SLA1 RNA and four uridines in the SL RNA exon are indicated by red circles.

(C) Predicted base-pairing interactions between vtRNA and the SL RNA 5′ splice site and branch-site sequence during SL *trans*-splicing.

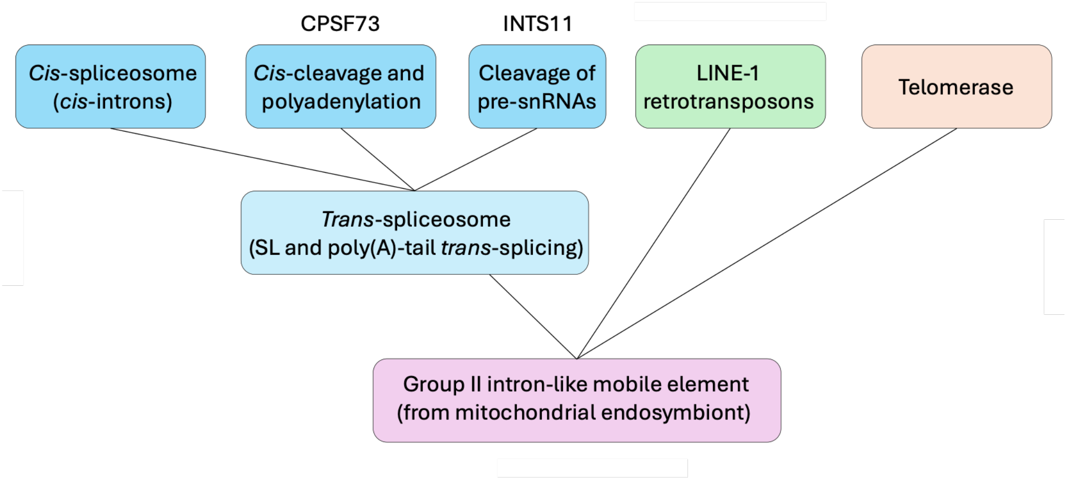


**Figure S10. Potential evolutionary links among *cis*- and *trans*-splicing, mRNA 3′-end formation, and snRNA 3′-end formation**
