## Supplementary material for "A *trans-*splicing model for poly(A)-tail addition during VSG mRNA maturation in *Trypanosoma brucei*": Instructions for original code

This script searches one or more FASTA or FASTQ files for a defined list of target nucleotide sequences and exports matching records to CSV files. It is intended to support reproducible sequence screening across macOS, Windows, and Linux.

**Requirements**

- Python 3.9 or later
- No external Python packages are required
- Input files must be plain-text FASTA or FASTQ files
- Compressed files such as .gz are not supported unless decompressed first

**Recommended Folder Structure**

Place the script, target-sequence file, and input files in a simple project folder.

```
sequence_search_project/
  batch_seq_search.py
  sequences.csv
  input/
    sample_1.fasta
    sample_2.fastq
  output/
```

The output/ folder can be empty or omitted before the first run. The script will create it if needed.

**Input Files**

**1. Script**

The main script should be named:

```
batch_seq_search.py
```

**2. Target Sequence File**

The target-sequence file should be named:

```
sequences.csv
```

It must contain two columns:

```
sequence_num,target_sequence
1,GGAGGTCATCCGACCC
2,GCTGCTACTGGGAGCTTCTC
3,TATAGTACAGAAACTGTTC
```

Required column meanings:

| Column | Description |
| --- | --- |
| `sequence_num` | Numeric identifier for the target sequence |
| `target_sequence` | Nucleotide sequence to search for |

The script performs exact substring matching. It does not perform alignment, approximate matching, mismatch tolerance, or reverse-complement searching unless those sequences are added explicitly to sequences.csv.

**3. Input Sequence Files**

Place FASTA or FASTQ files in the `input/` folder.

Supported file extensions:

```
.fasta
.fa
.fastq
.fq
```

The script searches files directly inside the input folder. It does not search nested subfolders.

Example FASTA input:

```
>read_001 sample sequence containing target 1
AAAGGAGGTCATCCGACCCTTT
>read_002 sample sequence with no target
AACCGGTTAACCGGTT
```

Example FASTQ input:

```
@read_001 sample sequence containing target 1
AAAGGAGGTCATCCGACCCTTT
+
IIIIIIIIIIIIIIIIIIIIII
```

**Installation**

**MacOS and Linux**

Open Terminal, navigate to the project folder, and confirm Python is available:

```
python3 --version
```

No package installation is required.

**Windows**

Open Command Prompt or PowerShell, navigate to the project folder, and confirm Python is available:

```
py --version
If py is unavailable, try:
python --version
```

No package installation is required.

**Running the Script**

**MacOS and Linux**

From the project folder:

```
python3 batch_seq_search.py --input-dir input_fasta --output-dir output_csv -
-sequences sequences.csv --progress-every 1000
```

**Windows**

From the project folder:

```
py batch_seq_search.py --input-dir input --output-dir output --sequences
sequences.csv –-progress every 1000
```

If using python instead of py:

```
python batch_seq_search.py --input-dir input --output-dir output --sequences
sequences.csv –-progress every 1000
```

For large datasets, the progress interval can be adjusted:

```
python3 batch_seq_search.py --input-dir input --output-dir output --sequences
sequences.csv --progress-every 10000
```

**Output Structure**

After the script runs, the output folder will contain:

```
output/
  summary.csv
  sample_1/
    summary.csv
    seq0001_sample_1_cleaned.csv
    seq0002_sample_1_cleaned.csv
    seq0003_sample_1_cleaned.csv
    ...
  sample_2/
```

```
summary.csv
seq0001_sample_2_cleaned.csv
seq0002_sample_2_cleaned.csv
seq0003_sample_2_cleaned.csv
...
```

**Output File Descriptions**

**Master Summary File**

`output/summary.csv`

This file summarizes all input files and all target sequences.

Columns:

| Column | Description |
|---|---|
| `file_name` | Input FASTA/FASTQ file name |
| `file_output_folder` | Output subfolder created for that input file |
| `sequence_num` | Target-sequence identifier from sequences.csv |
| `target_sequence` | Target sequence searched |
| `total_records_scanned` | Number of FASTA/FASTQ records scanned in the input file |
| `matches_found` | Number of records containing the target sequence |

Important: `matches_found` counts matching records, not the number of times a target appears. If a target appears multiple times in the same record, it is counted once for that record.

Example:

```
file_name,file_output_folder,sequence_num,target_sequence,total_records_scann
ed,matches_found
example_reads.fasta,example_reads,1,GGAGGTCATCCGACCC,4,2
example_reads.fasta,example_reads,2,GCTGCTACTGGGAGCTTCTC,4,1
example_reads.fasta,example_reads,3,TATAGTACAGAAACTGTTC,4,0
```

**Per-File Summary**

Each input file receives its own summary:

`output/sample_1/summary.csv`

This file has the same match-count information, limited to one input file.

Example:

```
file_name,sequence_num,target_sequence,total_records_scanned,matches_found
example_reads.fasta,1,GGAGGTCATCCGACCC,4,2
example_reads.fasta,2,GCTGCTACTGGGAGCTTCTC,4,1
example_reads.fasta,3,TATAGTACAGAAACTGTTC,4,0
```

**Per-Sequence Match Files**

For each target sequence, the script creates a CSV containing the records where that target was found.

Example file:

```
output/example_reads/seq0001_example_reads_cleaned.csv
```

Columns:

| Column | Description |
|---|---|
| `record_id` | First token from the FASTA/FASTQ record header |
| `description` | Remaining header text after the record ID |
| `sequence` | Full sequence from the matching record |

Example:

```
record_id,description,sequence
read_001,sample sequence containing target 1,AAAGGAGGTCATCCGACCCTTT
read_004,sample sequence containing target 1 again,TTTGGAGGTCATCCGACCCAAA
```

If no matches are found for a target, the corresponding CSV is still created but contains only the header row.
